# An Innovative Non-Hormonal Strategy Targeting Redox Active Metals to Down-Regulate Estrogen-, Progesterone-, Androgen- and Prolactin-Receptors in Breast Cancer

**DOI:** 10.1101/2023.02.02.526543

**Authors:** Faten Shehadeh-Tout, Heloisa H. Milioli, Suraya Roslan, Patric J. Jansson, Mahendiran Dharmasivam, Dinny Graham, Robin Anderson, Tharushi Wijesinghe, Mahan Gholam Azad, Des R. Richardson, Zaklina Kovacevic

## Abstract

Estrogen receptor-α (ER-α) is a key driver of breast cancer (BC) targeted by tamoxifen. However, tamoxifen resistance is a major problem. An important mechanism of resistance is the activation of EGFR/HER2/HER3 signaling and other hormone receptors (androgen receptor (AR), progesterone receptor (PR), prolactin receptor (PRL-R)) that intrinsically activate ER-α. Hence, therapeutics targeting multiple receptors, rather than ER-α alone, would be extremely useful and may overcome tamoxifen resistance. This study examined the activity of redox-active di-2-pyridylketone-4,4-dimethyl-3-thiosemicarbazone (Dp44mT) and di-2-pyridylketone-4-cyclohexyl-4-methyl-3-thiosemicarbazone (DpC), on the expression and activation of crucial hormone receptors, their co-factors, and key resistance pathways in ER-α-positive BC. Strikingly, DpC differentially regulated 106 estrogen-response genes with Sankey diagram analysis demonstrating this was linked to decreased mRNA levels of 4 central hormone receptors involved in BC pathogenesis, namely *ER*, *PR*, *AR*, and *PRL-R*. Mechanistic dissection demonstrated that due to DpC and Dp44mT binding metal ions, these agents caused a pronounced decrease in ER-α, AR, PR, and PRL-R protein expression. Ablation of the metal-binding site in the thiosemicarbazone totally prevented its suppressive activity, demonstrating a unique non-hormonal mechanism. DpC and Dp44mT also inhibited EGFR, HER2, and HER3 activation, their downstream signaling, and the expression of co-factors that promote ER-α transcriptional activity, including SRC3, NF-κB p65, and SP1. *In vivo,* DpC was highly tolerable and effectively inhibited ER-α-positive BC growth. In conclusion, through a bespoke non-hormonal mechanism targeting redox active metals, Dp44mT and DpC disrupt multiple key inter-receptor interactions between PR, AR, PRL-R, and tyrosine kinases that act with ER-α to promote BC, constituting an innovative therapeutic approach.

## Introduction

As the most common breast cancer (BC) type, estrogen receptor (ER)-positive BC is divided into two major sub-types, luminal A and luminal B, and is driven primarily by ER-α [1]. These sub-types are also characterized by the expression of other hormone receptors, namely progesterone receptor (PR; [2]), androgen receptor (AR; [3]), and prolactin receptor (PRL-R; [4]), which act solely, or in cooperation with ER-α to promote BC development and progression [5–9]. Hence, disrupting the interactions between these hormone receptors in BC, rather than inhibiting ER-α alone, may open a new frontier to successfully treat BCs and prevent disease relapse and metastasis [8].

Conventional treatments for ER-positive BC include agents that either block the synthesis of estrogen (*i.e.*, aromatase inhibitors; [10]), competitively bind to ER-α to inhibit its downstream signaling (*e.g.*, tamoxifen) [11], or bind and degrade ER-α (*e.g.*, Fulvestrant) [12]. Despite the availability of endocrine therapies and the good initial response to these agents, *de novo* and acquired resistance followed by relapse and metastasis is a common crisis for many BC patients [11]. Approximately 6-7% of new BC cases will have already metastasized when initially diagnosed, and up to 30% of early diagnosed, non-metastatic patients will eventually develop metastases [11].

Estradiol (E_2_)-binding enables phosphorylation at ER-α^Ser167^ and ER-α^Ser118^, leading to activation and nuclear translocation of ER-α [13]. In the nucleus, ER-α directly or indirectly binds to estrogen response elements (ERE) and promotes transcription of proliferation- and metastasis-related genes (*e.g.*, *collagenase*, *cyclin D*, *transforming growth factor-α (TGF-α)*, *c-myc*, *telomerase*, *etc.*) [14]. The ER-α also interacts with and binds to other proteins, including SRC1-3 and AP-1, that function as co-factors to initiate the transcription of other cancer-related genes [15]. In addition, ER-α also interacts with transcription factors, including SP1, c-Jun, and NF-кB p65, that promote and facilitate transcription, mainly by stabilizing ER-α-ERE binding [14]. This classical activation of ER-α *via* E_2_ and its subsequent binding to the ERE is known as the genomic signaling pathway [16].

The non-genomic signaling pathway of ER-α is a more rapid signaling cascade also initiated by the binding of E_2_ to ER-α and takes place in the cytoplasm [16]. This leads to the activation of mitogen-activated protein kinases (MAPKs) and phosphatidylinositol 3-kinase/protein kinase B (PI3K/AKT) to further drive cancer progression and metastasis [16]. This non-genomic signaling is facilitated by receptor tyrosine kinases (RTKs) such as human epidermal growth factor receptors (EGFR, HER2) and insulin growth factor-1 receptor (IGF-1R), which can interact with ER-α [16]. Importantly, these latter RTKs can also facilitate E_2_-independent activation of ER-α by phosphorylating it at multiple residues *e.g.*, ER-α^Ser-167^ and ER-α^Ser-118^ [13], which promotes endocrine resistance [10, 15, 16]. Thus, targeting the crosstalk between ER-α and RTKs may also constitute an effective way to treat BC and overcome endocrine resistance.

Potential, well-tolerated multi-focal therapies for BC are the redox-active thiosemicarbazones, di-2-pyridylketone-4,4-dimethyl-3-thiosemicarbazone (Dp44mT) and di-2-pyridylketone-4-cyclohexyl-4-methyl-3-thiosemicarbazone (DpC; [17–20]). The potent and selective anti-cancer activity of these compounds *in vitro* and *in vivo* involves a “double punch mechanism” involving the avid binding of tumor cell iron and copper that are critical for proliferation (first punch), followed by the formation of a redox-active iron or copper complex that generates cytotoxic reactive oxygen species (second punch) [18, 19, 21, 22]. These agents demonstrated marked safety *in vivo* and did not induce whole-body iron depletion due to the low doses required [18–20, 22]. A key characteristic of these ligands is their ability to up-regulate the metastasis suppressor, N-myc downstream-regulated gene 1 (NDRG1), *via* their ability to bind cancer cell iron [23–25], and this has been demonstrated to inhibit BC spread *in vivo* [26].

Additionally, increased expression of NDRG1 has been correlated with better BC prognosis and survival [27]. Both Dp44mT and DpC can effectively target and suppress multiple oncogenic signaling pathways in multiple tumor types, including the EGFR family (EGFR, HER2, HER3), NF-κB, PI3K/AKT, Wnt/TnC, YAP/TAZ, MAPK, *etc.* [28–31], all of which contribute to BC progression and resistance. Due to the marked and selective activity of these agents, the lead compound, DpC, entered Phase I clinical trials for advanced and resistant cancer [32]. Considering this, these novel agents could be a promising therapeutic strategy for BC with the current study examining the activity of Dp44mT and DpC on the expression of key drivers of this disease, including ER-α, PR, AR, and PRL-R. Additionally, the effects of DpC and Dp44mT were also investigated on major downstream signaling pathways and co-activators of ER-α in BC, with their efficacy being compared to the active metabolite of tamoxifen, 4-hydroxy-tamoxifen (4-OHT) [33].

Herein, we demonstrate the unexpected finding that through their ability to bind redox-active metal ions, Dp44mT and DpC markedly inhibited the protein expression of ER-α, PR, AR, and PRL-R in luminal-A and -B BC cell-types. Ablation of the metal-binding site of thiosemicarbazones totally blocked their activity. These agents also inhibited ER-α activation in response to E_2_, leading to the suppression of key downstream signaling pathways and effectively suppressing tumor growth *in vivo*, being very well tolerated. Further, the expression of multiple co-factors, namely SRC3, SP1, and JAK2, that promote ER-α activity was also decreased by these agents. This study is significant as the most common BC-type, ER-positive BC, is driven by ER-α and characterized by the interaction with PR, AR, PRL-R, and tyrosine kinases, which act integrally with ER-α to aggressively promote BC [5–9]. Thus, rather than inhibiting ER-α alone with tamoxifen, which leads to deadly resistance, the ability of Dp44mT and DpC to disrupt these multiple, key inter-receptor interactions, constitutes an innovative BC treatment strategy mediated *via* a novel non-hormonal mechanism targeting redox-active metals.

## Materials and Methods

### Cell culture

Luminal A human BC cells (MCF-7 and T47D) were a gift from Dr. M. Murray (The University of Sydney, NSW, Australia). Luminal B ZR-75-1 cells were obtained from Dr. D. Graham (The University of Sydney, NSW, Australia). MCF-7 and T47D cells were grown in Dulbecco’s modified eagle medium (DMEM), while ZR-75-1 cells were grown in Roswell Park Memorial Institute Medium 1640 (RPMI 1640) medium (Sigma-Aldrich, USA). DMEM and RPMI 1640 media were supplemented with 10% (v/v) fetal calf serum (Gibco, Australia), 1% (v/v) streptomycin/penicillin/glutamine, 1% (v/v) non-essential amino acids, and 1% (v/v) sodium pyruvate (Invitrogen, VIC, Australia). Insulin was used only for ZR-75-1 cells (0.25 U/mL). All supplements were obtained from Sigma-Aldrich.

### Spheroid culture

The 3-dimensional (3D) culture studies were performed using MCF-7 cells and low melting point agarose (Promega, Australia). Agarose was suspended in PBS at a concentration of 0.75% (w/v). Sterilized agarose (40 µL/well) was used to coat 96-well plates, with 100 µL of MCF-7 cells (3 x 10^5^ cells/mL) being seeded/well and incubated for 48 h/37°C. Cellular growth and viability were assessed by phase contrast microscopy and Trypan blue (Sigma-Aldrich) staining.

### Cell treatments

Cells were incubated for 24 h/37°C with control medium or medium containing either Desferrioxamine (DFO; 250 μM; Novaris), 4-OHT (5 µM; Sigma-Aldrich), Dp44mT (5 μM), DpC (5 μM), or the negative control compound, 2-benzoylpyridine-2-methyl-3-thiosemicarbazone (Bp2mT; 5 μM), which was specifically designed not to bind metal ions [17, 34–36]. The agents, Dp44mT, DpC, and Bp2mT were synthesized and characterized, as described previously [37, 38]. For E_2_ treatment, T47D and MCF-7 cells were incubated in phenol red-free charcoal-stripped medium with E_2_ (10 nM; Sigma-Aldrich) for 5 min and 15 min/37°C, respectively. For proteasomal inhibition studies, T47D and MCF-7 cells were treated with proteasomal inhibitor MG132 (Sigma-Aldrich) at 2.5 and 5 µM, respectively, in the presence of Dp44mT or DpC for 24 h/37°C.

### RNA isolation, sequencing, and analysis

MCF-7 cells were treated with either control medium or medium containing DpC (5 μM) for 24 h/37°C. Isolation of mRNA was then performed using TRIzol^®^ (Invitrogen) according to the manufacturers’ procedures. RNA sequencing (RNAseq) was performed by next-generation sequencing (NGS) using Illumina TruSeq technology (Ramaciotti Centre for Genomics, University of New South Wales, Sydney, Australia). Sequences were trimmed using Trim Galore (version 0.4.4; Babraham Institute UK), and genome-guided alignment to a human reference (HG38) was performed using STAR software (version 2.5; National Human Genome Research Institute, USA). Duplicated reads were located and tagged using the Picard tool, MarkDuplicates (version 1.138; Broad Institute, USA). Differential gene expression was computed in R (version 3.4.3; Lucent Technologies, USA) using the package, Rsubread 1.28.0 [39]. Differential gene expression analysis was conducted using the edgeR 3.20.1 [40] and limma 3.34.1 [41] packages (Bioconductor, USA). Gene set enrichment analysis (GSEA) was done with clusterProfiler. Data from RNAseq were deposited in the GEO repository (GEO accession number, GSE192942).

### Western blot

Cell lysate preparation and immunoblotting was performed according to established protocols [29]. The primary antibodies included the following from Cell Signaling Technology, MA, USA: ER-α (Cat. #:13258), AR (Cat. #:5153s), PR (Cat. #:8757s), PRL-R (Cat. #:13552s), p-ER-α^Ser167^ (Cat. #: 64508), p-ER-α^Ser 118^ (Cat. #:2511), c-Fos (Cat. #:2250s), p-c-Fos^Ser32^ (Cat. #: 5348s), p53 (Cat. #: 48818s), SRC3 (Cat. #: 5765s), c-SRC (Cat. #: 2109s), p-c-SRC^Tyr416^ (Cat. #: 6943s), c-Jun (Cat. #:9165s), p-c-Jun^Ser63^ (Cat. #: 2361s), MDM2 (Cat. #: 86934s), p-MDM2^Ser166^ (Cat. #: 3521s) SP1 (Cat. #:9389s), JAK2 (Cat. #:3230), PRL-R (Cat. #: 13552s), NF-κB p65 (Cat. #: 8242s), HER2 (Cat. #:4290s), p-HER2^Tyr1221/1222^ (Cat. #: 2243s), HER3 (Cat. #:12708s), EGFR (Cat. #:4267s), p-EGFR^Tyr1086^ (Cat. #: 2220s), p-EGFR^Tyr1068^ (Cat. #:2234s), IGF-IRβ (Cat. #: 3027s), p-IGF-^IRβTyr 1135^ (Cat. #: 3918s), P38-MAPK (Cat. #: 9212), ERK1/2 (Cat. #:9107), p-ERK1^Thr202^/p-ERK2^Tyr204^ (Cat. #: 4370, MEK1 (Cat. #:9124s), MEK1 (Cat. #:9125s), p-MEK^Ser217/221^ (Cat. #:9121s), AKT (Cat. #:9272s), p-AKT^Thr308^ (Cat. #:9275s), p-AKT^Ser473^ (Cat. #:9271s), mTOR (Cat. #:2983s), PI3K-p110 (Cat. #:3011s), PI3K-p85 (Cat. #:4292s), and p-PI3K p85^Tyr458^/p55^Tyr199^ (Cat. #:4228s). NDRG1 antibody was from Abcam, UK (Cat. #: ab37897). The secondary antibodies utilized were from Sigma-Aldrich and included anti-mouse (Cat. #: A4416), anti-rabbit (Cat. #: A6154), and anti-goat (Cat. #: A5420), and these were prepared at a 1:10,000 dilution. As a loading control, an anti-β-actin antibody (1:10,000; Sigma-Aldrich) was used.

### RT-PCR

RT-PCR was performed according to standard procedures [29] using the following primers: ER-α (F: 5ꞌ-GAAGAGGAGGGAGAATGTTG-3ꞌ; R: 5ꞌ-ACTGAAGGGTCTGGTAGGAT-3ꞌ); PR (F: 5ꞌ-AATGGTGTTTGGTCTAGGATG-3ꞌ; R: 5ꞌ-TGTAAGTTGATAGAAACGCTGT-3ꞌ); AR (F: 5ꞌ-CCCTACGGCTACACTCGG-3ꞌ; R: 5ꞌ-AGGTGCCTCATTCGGACA-3ꞌ); PRL-R (F: 5ꞌ-TCATGATGGTCAATGCCACT-3ꞌ; R: 5ꞌ-GCGTGAACCAACCAGTTTTT-3ꞌ); and β-actin (F: 5ꞌ-CCCGCCGCCAGCTCACCATGG-3ꞌ; R: 5ꞌ-AAGGTCTCAAACATGATCTGGGTC-3ꞌ).

### Small interfering RNA and gene silencing

Gene silencing was performed following the manufacturer’s protocol. Briefly, at 75-85% confluence, cells were transfected with 20 nM select siRNA for either c-Jun (si-c-Jun; Sigma-Aldrich), p53 (si-p53; Sigma-Aldrich), 50 nM select siRNA for NDRG1 (siNDRG1), or a negative control siRNA (si-Con; Sigma-Aldrich) using Lipofectamine RNAmax (ThermoFisher). Cells were incubated for 48 h/37°C prior to protein extraction. To examine the effect of Dp44mT and DpC on silenced cells, siRNAs were incubated with cells for 24 h/37°C, followed by incubation with Dp44mT and DpC treatment for a further 24 h/37°C.

### Immunofluorescence

Immunofluorescence was carried out following standard procedures [29]. Slides were incubated overnight with primary antibodies against ER-α (Cat. #: 13258; 1:500), AR (Cat. #: 5153; 1:400), or PR (Cat. #: 8757; 1:400) diluted in 1% BSA dissolved in PBS/4°C. After removing the primary antibodies, cells were washed and incubated with secondary antibodies (anti-mouse Cat. # A-21200 and anti-rabbit Cat. # A-11012; Invitrogen) at a dilution of 1:1000 for 1 h/20°C.

Spheroids were treated with control medium or medium containing Dp44mT (5 µM) or DpC (5 µM) for 24 h/37°C. Spheroids (10-20) were then collected in an Eppendorf tube, and immunofluorescence performed using standard procedures [29]. Briefly, spheroids were washed, fixed using 4% paraformaldehyde (Sigma-Aldrich) for 1 h/20°C, permeabilized with Triton-X (1%) for 1 h/20°C, blocked with 5% BSA overnight at 4°C, and incubated with primary antibodies overnight at 4°C. Spheroids were then incubated with anti-rabbit or anti-mouse antibodies (1:1000; Cat #. A-21200 and Cat #. A-11012; Invitrogen) and fixed in 1% agarose using Prolong Gold Antifade mounting solution containing DAPI (ThermoFisher) for 15 min/20°C. The spheroids were imaged using a ZEISS LSM 800 plus Airyscan laser confocal microscope (10×; Zeiss, Jena, Germany), and the images analyzed using ImageJ software (NIH, USA).

### *In vivo* studies

Briefly, NOD-SCID-Gamma (NSG) mice (6–8 weeks; Australian BioResources) were subcutaneously implanted with E_2_ 90-day slow-release pellets (20 ng/day; Belma Technologies, Belgium) four days prior to injection of cancer cells. The orthotopic xenograft was established using MCF-7 (2 x 10^7^) cells that were injected into the fat pads of NSG female mice. Once palpable tumors of 200 mm^3^ formed 23 days post-implantation of the cancer cells, mice were randomized into 3 groups (*n* = 10), with each group receiving 100 µL of either DpC (5 mg/kg, 3x/week; oral gavage, dissolved in propylene glycol), tamoxifen (35 mg, 60-day slow-release pellet implanted intradermally; Innovative Research of America, USA), or the vehicle control (30% propylene glycol in saline given orally) for another 29 days. Major organs were harvested and fixed with 4% PFA for a least 24 h/20 °C. All animal procedures were conducted with approval from Austin Health Animal Ethics Committee, Melbourne.

### Immunohistochemistry

Slides were prepared and stained, as described [17]. The primary antibodies, ER-α (1:200, Cat #: MA5-14501, Invitrogen), AR (1:300; Cat #: 5153; Cell Signaling Technology), and PR (1:700; Cat #: 8757; Cell Signaling Technology) were used to detect these proteins in formalin-fixed, paraffin-embedded tumor specimens.

### Evaluation of immunostaining

ImageJ was used to quantify DAB staining. For each image, the intensity was converted to an optical density (OD) using the formula: OD = log (max intensity/mean intensity). These values were averaged across samples in each treatment group to give a mean DAB intensity.

### Densitometry and statistics

Densitometry was performed using Image Lab software (Bio-Rad, USA) and Quantity One v4.6.9 (Bio-Rad, USA) to quantify protein and mRNA, respectively. Data were normalized using the relative β-actin loading control. All blots are typical of three independent experiments and are presented as the mean ± standard deviation (S.D). Results were compared using the Student’s *t*-test and considered statistically significant when *p* < 0.05.

## Results

### DpC down-regulates estrogen response genes, while increasing markers of apoptosis and activation of the p53 signaling pathway in MCF-7 BC cells

We initially assessed the effect of our lead agent, DpC (5 µM), on MCF-7 cells after a 24 h/37°C treatment relative to the vehicle-treated control by performing whole-genome RNAseq (**Fig. 1**). The mRNA expression analysis revealed 1521 genes with a significant differential in levels induced by DpC relative to the control (**Supplemental Table 1**). KEGG pathway analysis demonstrated in DpC-treated cells *versus* control cells that there was marked up-regulation of MAPK, hypoxia-inducible factor-1 (HIF-1), apoptosis, p53, and mammalian target of rapamycin (mTOR) signaling, as well as increased protein processing in the endoplasmic reticulum (**Fig. 1A**). Endocrine resistance and steroid biogenesis pathways were also significantly increased, while *N*-glycan biosynthesis and the estrogen signaling pathways were potently decreased in response to DpC treatment *versus* the control (**Fig. 1A**).

**Figure 1.**
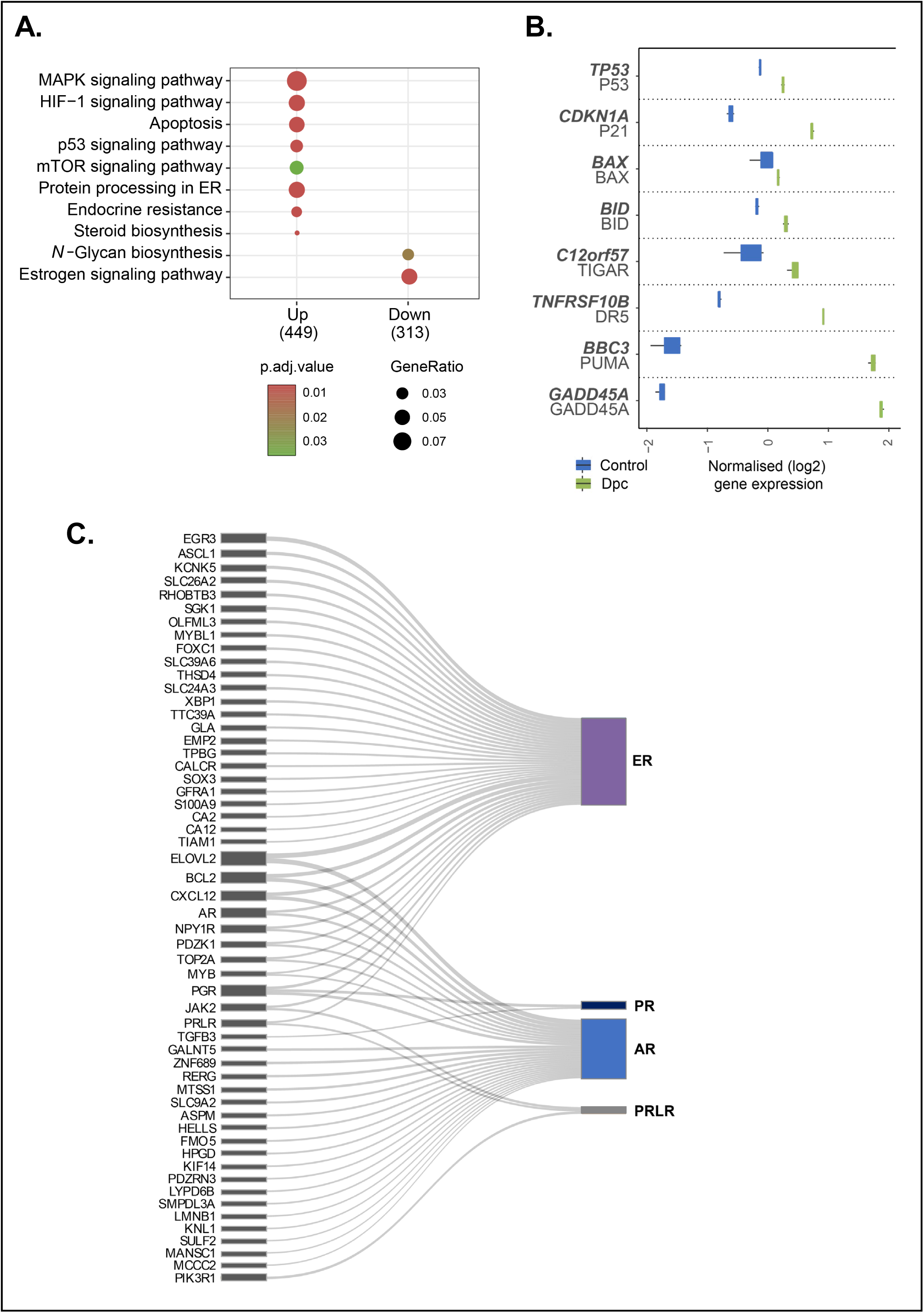
RNAseq analysis of gene expression affected by DpC in MCF-7 cells. MCF-7 cells were incubated with control medium or this medium containing DpC (5 µM) for 24 h, and RNA sequencing was performed. **(A)** KEGG pathway analysis of the top up- and down-regulated signaling pathways. **(B)** Expression of p53-regulated and apoptosis-inducing genes. **(C)** Sankey diagram analysis of RNAseq data demonstrating genes differentially regulated by DpC linked to either ER, PR, AR, or the PRL-R.

The increased p53 signaling and apoptosis gene expression signature after incubation with DpC were further confirmed by examining mRNAs associated with p53 regulation and apoptosis (**Fig. 1B**). This analysis demonstrated that relative to the control, DpC significantly up-regulated genes encoding p53, p21, BCL2 associated X (BAX), BH3 interacting domain death agonist (BID), TP53-induced glycolysis regulatory phosphatase (TIGAR), death receptor 5 (DR5), p53 up-regulated modulator of apoptosis (PUMA), and growth arrest and DNA damage-inducible alpha (GADD45A; **Fig. 1B**).

The top 35 most significantly up-regulated genes in response to DpC relative to the control include the well-established target of this agent, namely the metastasis suppressor, N-myc downstream-regulated gene 1 (*NDRG1*; [25]), an inhibitor of BC metastasis (*KLF6*; [42]), and the growth arrest gene, *GADD45A* (**Table 1**). The top 35 genes down-regulated by DpC include key hormone receptors (*i.e., estrogen receptor 1* (*ESR1*)*, progesterone receptor (PGR)*), metallopeptidases (*i.e., ADAMTS9, ADAMTS19*), and key proliferation genes (*i.e., EGR3, CXCL12, SOX2*; **Table 2**).

**Table 1:** Top 35 most significantly up-regulated genes in response to DpC-treated vs. vehicle control-treated MCF-7 cells. The table shows log fold change (logFC), average expression across all samples, in log2 counts per million reads (AveExpr), logFC divided by its standard error (t), raw *p*-value (P.Value) and Benjamini-Hochberg false discovery rate adjusted *p*-value (adj.P.Val).

| ENTREZ ID | SYMBOL | MAP | logFC | AveExpr | t | P.Value | adj.P.Val |
| --- | --- | --- | --- | --- | --- | --- | --- |
| 3099 | HK2 | 2p12 | 4.33 | 8.66 | 91.31 | 7.83E-15 | 7.91E-11 |
| 8974 | P4HA2 | 5q31.1 | 4.17 | 8.46 | 77.41 | 3.39E-14 | 1.71E-10 |
| 664 | BNIP3 | 10q26.3 | 4.39 | 8.05 | 67.68 | 1.12E-13 | 2.17E-10 |
| 54583 | EGLN1 | 1q42.2 | 5.38 | 8.50 | 66.50 | 1.34E-13 | 2.17E-10 |
| 23645 | PPP1R15A | 19q13.33 | 4.23 | 7.79 | 64.92 | 1.62E-13 | 2.17E-10 |
| 230 | ALDOC | 17q11.2 | 5.62 | 7.08 | 64.91 | 1.67E-13 | 2.17E-10 |
| 2026 | ENO2 | 12p13.31 | 4.56 | 7.41 | 64.59 | 1.71E-13 | 2.17E-10 |
| 10397 | NDRG1 | 8q24.22 | 6.32 | 8.74 | 64.78 | 1.73E-13 | 2.17E-10 |
| 7128 | TNFAIP3 | 6q23.3 | 4.44 | 6.69 | 63.50 | 1.98E-13 | 2.17E-10 |
| 467 | ATF3 | 1q32.3 | 7.44 | 5.59 | 62.78 | 2.35E-13 | 2.17E-10 |
| 7779 | SLC30A1 | 1q32.3 | 5.44 | 7.84 | 62.37 | 2.37E-13 | 2.17E-10 |
| 1316 | KLF6 | 10p15.2 | 3.43 | 7.82 | 60.82 | 2.87E-13 | 2.37E-10 |
| 55818 | KDM3A | 2p11.2 | 3.91 | 8.91 | 60.44 | 3.04E-13 | 2.37E-10 |
| 5163 | PDK1 | 2q31.1 | 4.51 | 6.90 | 57.30 | 4.93E-13 | 3.56E-10 |
| 4189 | DNAJB9 | 7q31.1 | 4.51 | 7.60 | 54.99 | 7.11E-13 | 4.79E-10 |
| 8537 | BCAS1 | 20q13.2 | 2.83 | 8.57 | 53.64 | 8.72E-13 | 5.50E-10 |
| 30001 | ERO1A | 14q22.1 | 4.12 | 9.61 | 52.91 | 9.95E-13 | 5.91E-10 |
| 5236 | PGM1 | 1p31.3 | 3.76 | 6.21 | 52.08 | 1.14E-12 | 6.40E-10 |
| 8553 | BHLHE40 | 3p26.1 | 3.50 | 9.01 | 51.63 | 1.23E-12 | 6.53E-10 |
| 5230 | PGK1 | Xq21.1 | 3.45 | 10.71 | 51.16 | 1.33E-12 | 6.73E-10 |
| 7422 | VEGFA | 6p21.1 | 3.95 | 8.31 | 50.21 | 1.58E-12 | 7.60E-10 |
| 494470 | RNF165 | 18q21.1 | 6.93 | 5.43 | 49.13 | 2.04E-12 | 9.38E-10 |
| 284716 | RIMKLA | 1p34.2 | 4.08 | 5.73 | 47.80 | 2.45E-12 | 1.07E-09 |
| 5210 | PFKFB4 | 3p21.31 | 6.15 | 7.36 | 47.76 | 2.57E-12 | 1.07E-09 |
| 4490 | MT1B | 16q13 | 13.05 | 2.42 | 48.56 | 2.64E-12 | 1.07E-09 |
| 7045 | TGFBI | 5q31.1 | 3.03 | 7.47 | 47.05 | 2.79E-12 | 1.07E-09 |
| 3486 | IGFBP3 | 7p12.3 | 3.25 | 7.37 | 46.86 | 2.90E-12 | 1.07E-09 |
| 5209 | PFKFB3 | 10p15.1 | 3.69 | 9.48 | 46.78 | 2.95E-12 | 1.07E-09 |
| 9819 | TSC22D2 | 3q25.1 | 2.60 | 7.53 | 46.04 | 3.38E-12 | 1.11E-09 |
| 2997 | GYS1 | 19q13.33 | 4.16 | 8.65 | 46.09 | 3.39E-12 | 1.11E-09 |
| 1847 | DUSP5 | 10q25.2 | 3.28 | 5.14 | 45.95 | 3.45E-12 | 1.11E-09 |
| 1647 | GADD45A | 1p31.3 | 3.82 | 5.58 | 45.88 | 3.51E-12 | 1.11E-09 |
| 84560 | MT4 | 16q13 | 9.60 | 0.06 | 46.18 | 3.80E-12 | 1.16E-09 |
| 29923 | HILPDA | 7q32.1 | 2.64 | 6.96 | 44.37 | 4.69E-12 | 1.39E-09 |
| 9709 | HERPUD1 | 16q13 | 2.74 | 6.78 | 43.33 | 5.78E-12 | 1.67E-09 |

**Table 2:** Top 35 most significantly down-regulated genes in response to DpC-treated vs. vehicle control-treated MCF-7 cells. The table shows log fold change (logFC), average expression across all samples, in log2 counts *per* million reads (AveExpr), logFC divided by its standard error (t), raw p-value (P.Value) and Benjamini-Hochberg false discovery rate adjusted *p*-value (adj.P.Val).

| ENTREZ ID | SYMBOL | MAP | logFC | AveExpr | t | P.Value | adj.P.Val |
| --- | --- | --- | --- | --- | --- | --- | --- |
| 2099 | ESR1 | 6q25.1-q25.2 | -2.90 | 6.74 | -42.20 | 7.33E-12 | 1.95E-09 |
| 1960 | EGR3 | 8p21.3 | -3.85 | 4.06 | -37.83 | 1.94E-11 | 4.26E-09 |
| 5087 | PBX1 | 1q23.3 | -3.09 | 4.83 | -36.42 | 2.70E-11 | 5.33E-09 |
| 341640 | FREM2 | 13q13.3 | -3.54 | 5.57 | -35.40 | 3.48E-11 | 6.51E-09 |
| 118980 | SFXN2 | 10q24.32 | -2.60 | 5.67 | -33.99 | 4.97E-11 | 8.03E-09 |
| 23741 | EID1 | 15q21.1 | -2.38 | 6.92 | -30.64 | 1.24E-10 | 1.57E-08 |
| 54898 | ELOVL2 | 6p24.2 | -4.25 | 3.54 | -30.50 | 1.31E-10 | 1.61E-08 |
| 6387 | CXCL12 | 10q11.21 | -3.14 | 3.79 | -29.92 | 1.53E-10 | 1.74E-08 |
| 8503 | PIK3R3 | 1p34.1 | -3.49 | 5.75 | -29.89 | 1.55E-10 | 1.74E-08 |
| 51473 | DCDC2 | 6p22.3 | -2.81 | 5.46 | -28.81 | 2.14E-10 | 2.14E-08 |
| 124540 | MSI2 | 17q22 | -2.13 | 6.60 | -28.67 | 2.23E-10 | 2.16E-08 |
| 135398 | C6orf141 | 6p12.3 | -2.22 | 6.35 | -28.52 | 2.33E-10 | 2.25E-08 |
| 219623 | TMEM26 | 10q21.2 | -3.63 | 2.78 | -27.20 | 3.56E-10 | 3.10E-08 |
| 55170 | PRMT6 | 1p13.3 | -2.53 | 6.12 | -26.52 | 4.43E-10 | 3.70E-08 |
| 1836 | SLC26A2 | 5q32 | -2.66 | 4.27 | -26.04 | 5.21E-10 | 4.12E-08 |
| 22845 | DOLK | 9q34.11 | -2.92 | 3.50 | -25.96 | 5.34E-10 | 4.18E-08 |
| 171019 | ADAMTS19 | 5q23.3 | -2.72 | 5.27 | -25.88 | 5.50E-10 | 4.18E-08 |
| 56999 | ADAMTS9 | 3p14.1 | -3.74 | 3.11 | -25.44 | 6.42E-10 | 4.74E-08 |
| 6657 | SOX2 | 3q26.33 | -4.10 | 2.70 | -25.30 | 6.78E-10 | 4.96E-08 |
| 105373957 | LOC105373957 | 2q37.3 | -3.18 | 2.48 | -25.18 | 7.01E-10 | 5.09E-08 |
| 10376 | TUBA1B | 12q13.12 | -2.01 | 8.29 | -24.86 | 7.82E-10 | 5.52E-08 |
| 79624 | ARMT1 | 6q25.1 | -2.59 | 6.68 | -24.66 | 8.39E-10 | 5.80E-08 |
| 9630 | GNA14 | 9q21.2 | -3.20 | 3.01 | -24.34 | 9.43E-10 | 6.48E-08 |
| 55250 | ELP2 | 18q12.2 | -2.04 | 7.56 | -23.97 | 1.08E-09 | 7.31E-08 |
| 7494 | XBP1 | 22q12.1 | -1.98 | 9.37 | -23.74 | 1.17E-09 | 7.78E-08 |
| 399694 | SHC4 | 15q21.1 | -2.71 | 3.26 | -23.73 | 1.18E-09 | 7.78E-08 |
| 22836 | RHOBTB3 | 5q15 | -2.34 | 7.13 | -23.45 | 1.31E-09 | 8.56E-08 |
| 338324 | S100A7A | 1q21.3 | -4.18 | 0.56 | -23.42 | 1.34E-09 | 8.62E-08 |
| 7026 | NR2F2 | 15q26.2 | -2.42 | 6.78 | -23.02 | 1.53E-09 | 9.42E-08 |
| 25800 | SLC39A6 | 18q12.2 | -2.12 | 10.34 | -22.40 | 1.95E-09 | 1.15E-07 |
| 22996 | TTC39A | 1p32.3 | -1.94 | 5.59 | -22.32 | 2.01E-09 | 1.18E-07 |
| 93183 | PIGM | 1q23.2 | -3.18 | 2.75 | -22.29 | 2.04E-09 | 1.19E-07 |
| 101927457 | LINC01522 | 20q13.13 | -4.54 | 1.57 | -22.19 | 2.16E-09 | 1.25E-07 |
| 5241 | PGR | 11q22.1 | -2.37 | 4.08 | -22.13 | 2.17E-09 | 1.25E-07 |
| 283310 | OTOGL | 12q21.31 | -4.24 | 0.53 | -22.14 | 2.19E-09 | 1.26E-07 |

Since the current focus was estrogen-positive MCF-7 BC cells, studies further examined the effect of DpC on the hallmark estrogen-response gene set. Surprisingly, DpC differentially regulated 106 estrogen-response genes when compared to the control, demonstrating its broad effect on estrogen signaling (**Tables 3, 4**). A number of E_2_-mediated genes related to inhibition of growth were up-regulated by DpC in MCF-7 cells, including *CCNA1* and *FOS* (**Table 3**). In addition, there was down-regulation of hormone receptors (*i.e., PGR* and *PRL-R*), co-activators of hormone receptors (*i.e., ASCL1, MYB,* and *SOX3*), transcription factors (*i.e., FOXC1*), anti-apoptosis genes (*BCL2)*, and tyrosine kinases (*i.e., JAK2;* **Table 4**). Using a Sankey diagram, the differentially expressed gene set demonstrated an intertwined link with 4 major hormone receptors involved in BC pathogenesis, namely ER, PR, AR, and PRL-R [5–9] (**Fig. 1C**). These results suggested that DpC is responsible for a broad inhibitory effect on hormone signaling in BC.

**Table 3:** Estrogen response genes up-regulated in response to DpC-treated *vs.* vehicle control-treated MCF-7 cells. The table shows log fold change (logFC), average expression across all samples, in log2 counts *per* million reads (AveExpr), logFC divided by its standard error (t), raw *p*-value (P.Value) and Benjamini-Hochberg false discovery rate adjusted *p*-value (adj.P.Val).

| ENTREZ ID | SYMBOL | MAP | logFC | AveExpr | t | P.Value | adj.P.Val |
| --- | --- | --- | --- | --- | --- | --- | --- |
| 23650 | TRIM29 | 11q23.3 | 4.33 | 5.35 | 31.88 | 8.9E-11 | 1.3E-08 |
| 3918 | LAMC2 | 1q25.3 | 2.56 | 6.10 | 21.70 | 2.6E-09 | 1.4E-07 |
| 3669 | ISG20 | 15q26.1 | 3.08 | 5.60 | 11.78 | 5.0E-07 | 9.8E-06 |
| 2288 | FKBP4 | 12p13.33 | 1.76 | 10.62 | 11.65 | 5.5E-07 | 1.0E-05 |
| 8839 | WISP2 | 20q13.12 | 2.95 | 3.30 | 10.44 | 1.4E-06 | 2.3E-05 |
| 8140 | SLC7A5 | 16q24.2 | 2.43 | 8.87 | 7.81 | 1.4E-05 | 1.7E-04 |
| 2353 | FOS | 14q24.3 | 1.81 | 6.04 | 7.22 | 2.6E-05 | 3.0E-04 |
| 6566 | SLC16A1 | 1p13.2 | 1.37 | 4.48 | 6.09 | 9.5E-05 | 9.5E-04 |
| 8900 | CCNA1 | 13q13.3 | 1.69 | 3.10 | 6.09 | 9.5E-05 | 9.6E-04 |
| 2222 | FDFT1 | 8p23.1 | 1.32 | 8.14 | 6.01 | 1.0E-04 | 1.0E-03 |
| 7869 | SEMA3B | 3p21.31 | 2.19 | 5.94 | 5.54 | 1.9E-04 | 1.8E-03 |
| 1152 | CKB | 14q32.33 | 2.25 | 5.64 | 4.46 | 8.1E-04 | 6.5E-03 |
| 26353 | HSPB8 | 12q24.23 | 1.31 | 5.83 | 4.39 | 8.9E-04 | 7.2E-03 |
| 1717 | DHCR7 | 11q13.4 | 1.42 | 8.54 | 4.17 | 1.2E-03 | 9.6E-03 |
| 2444 | FRK | 6q22.1 | 1.20 | 3.73 | 3.49 | 3.4E-03 | 2.5E-02 |
| 9314 | KLF4 | 9q31.2 | 1.13 | 7.33 | 2.58 | 1.5E-02 | 9.4E-02 |
| 4665 | NAB2 | 12q13.3 | 1.29 | 7.42 | 2.55 | 1.6E-02 | 9.8E-02 |
| 2810 | SFN | 1p36.11 | 1.49 | 6.23 | 1.81 | 5.2E-02 | 2.9E-01 |
| 51599 | LSR | 19q13.12 | 1.40 | 7.37 | 1.68 | 6.3E-02 | 3.5E-01 |
| 645 | BLVRB | 19q13.2 | 1.38 | 6.14 | 1.59 | 7.3E-02 | 4.0E-01 |
| 10202 | DHRS2 | 14q11.2 | 1.11 | 5.13 | 1.53 | 8.0E-02 | 4.4E-01 |
| 51195 | RAPGEFL1 | 17q21.1 | 1.09 | 7.16 | 1.35 | 1.0E-01 | 5.5E-01 |
| 11145 | PLA2G16 | 11q12.3-q13.1 | 1.09 | 4.26 | 1.11 | 1.5E-01 | 7.6E-01 |
| 9022 | CLIC3 | 9q34.3 | 1.52 | 4.22 | 1.11 | 1.5E-01 | 7.6E-01 |
| 7466 | WFS1 | 4p16.1 | 1.16 | 6.28 | 0.86 | 2.1E-01 | 1.0E+00 |
| 7033 | TFF3 | 21q22.3 | 1.09 | 4.92 | 0.34 | 3.7E-01 | 1.0E+00 |
| 11187 | PKP3 | 11p15.5 | 1.06 | 5.57 | 0.19 | 4.3E-01 | 1.0E+00 |

**Table 4:**
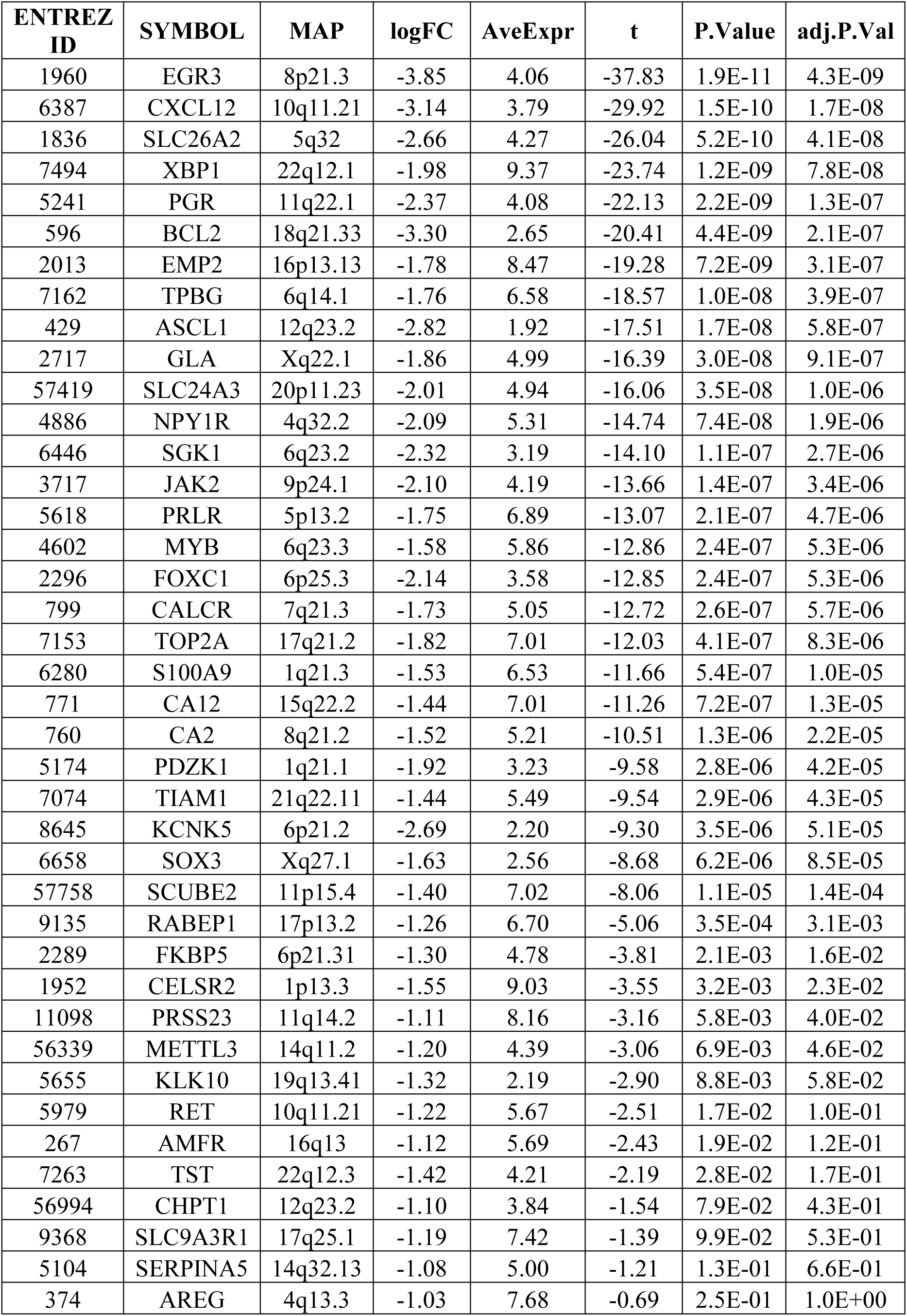
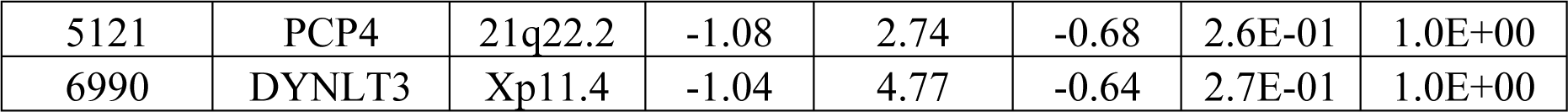
Estrogen response genes down-regulated in response to DpC-treated vs. vehicle control-treated MCF-7 cells. The table shows log fold change (logFC), average expression across all samples, in log2 counts per million reads (AveExpr), logFC divided by its standard error (t), raw *p*-value (P.Value) and Benjamini-Hochberg false discovery rate adjusted *p*-value (adj.P.Val).

Considering the major roles of the hormone receptors ER, AR, PR, and PRL-R and their interactions in BC progression [5–9], these findings provided a strong rationale for further investigating the effects of DpC on receptor protein levels and activation in ER-α-positive BC cells.

### Dp44mT and DpC markedly down-regulate ER-α, AR, PR, and PRL-R protein levels due to their metal-binding activity

Based on the results in **Figure 1**, studies next assessed the effects of Dp44mT (5 µM) and DpC (5 µM) on the protein expression of ER-α, PR, AR, and PRL-R in luminal A MCF-7 and T47D BC cells, as well as luminal B, ZR-75-1 cells. The key target of these thiosemicarbazones is their ability to bind iron and copper that leads to redox active complexes [17–20]. As such, Bp2mT (5 µM) was used as a negative control since it is a specially designed structural analog of Dp44mT and DpC that cannot bind metals [17, 34–36]. Further, the well-characterized, iron chelator, desferrioxamine (DFO; 250 µM), was used as a positive control, as this clinically used drug binds cellular iron pools, but does not form redox-active complexes like Dp44mT and DpC [43]. The higher concentration of DFO implemented is due to the low membrane permeability of this hydrophilic drug [43]. To compare the effects of Dp44mT and DpC to conventional BC endocrine therapies, experiments also examined the active metabolite of tamoxifen, namely 4-OHT (5 µM) [33, 44]. The cells were incubated with all agents for 24 h/37°C.

We initially examined the effects of these agents on ER-α in MCF-7 cells. Four major ER-α bands were identified at 36, 46, 56, and 66 kDa (**Fig. 2A**), which corresponds to its well-characterized protein isoforms [45] (**Fig. 2A**), with the 66 kDa band being the predominant isoform and the major driver of hormone-positive BC cells [45]. The 66 kDa band was quantified by densitometric analysis, as it is the focus of our investigation. Both Dp44mT and DpC caused a pronounced and significant reduction in the expression of the 66 kDa ER-α isoform, with the other 3 isoforms also being robustly decreased (**Fig. 2A**). Notably, 4-OHT significantly up-regulated ER-α relative to the control, which occurs due to the stabilization of ER-α and inhibition of its degradation when bound to 4-OHT [46].

**Figure 2.**
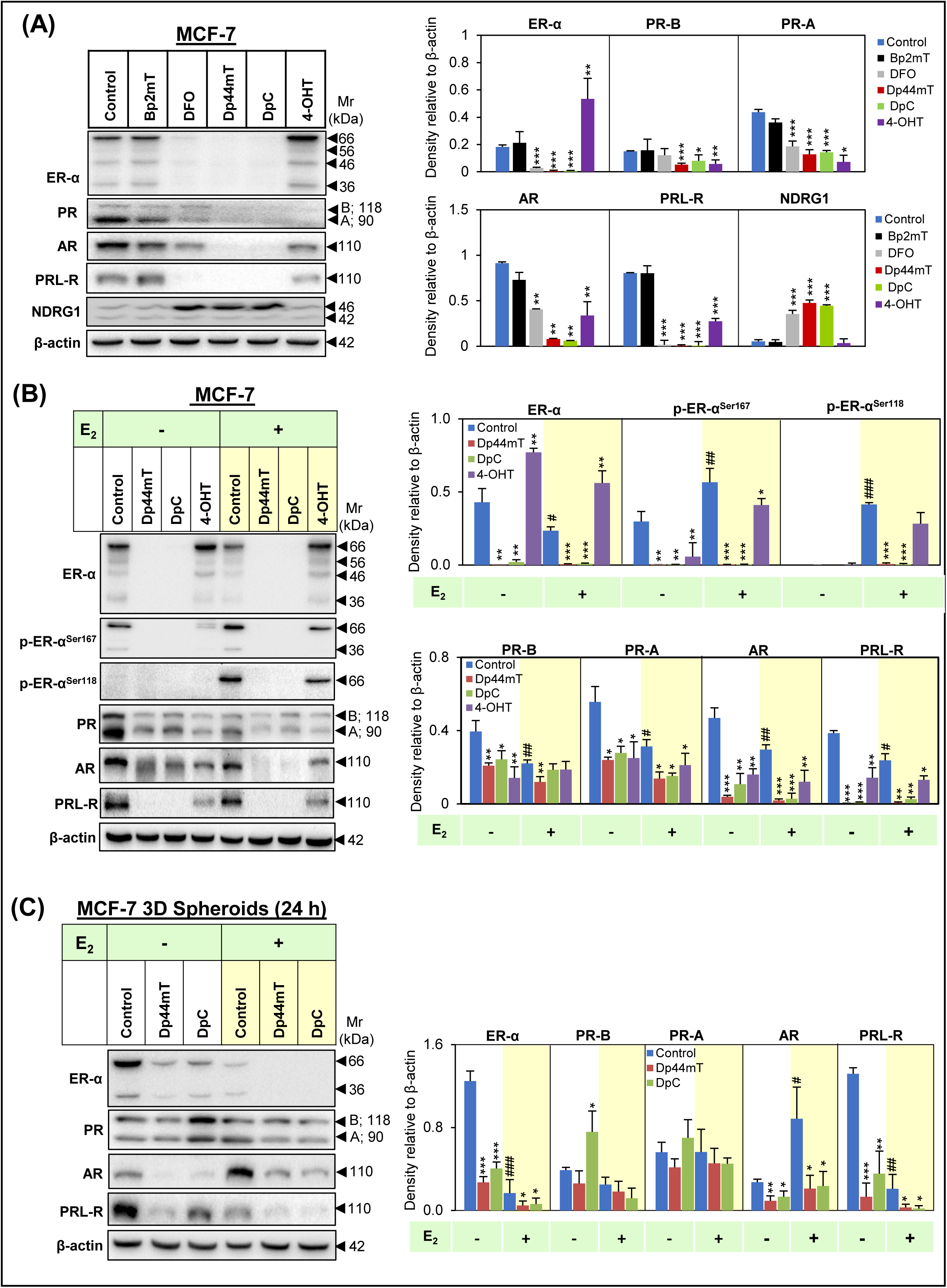
Dp44mT and DpC down-regulate ER-α, PR, AR, and PRL-R protein levels in MCF-7 cells, but up-regulate NDRG1. **(A)** MCF-7 cells were incubated with either control medium or this medium containing Bp2mT (5 µM), DFO (250 µM), Dp44mT (5 µM), DpC (5 µM), or 4-OHT (5 µM) for 24 h. Cellular lysates were then assessed for protein expression of the ER-α, PR, AR, PRL-R, and NDRG1 *via* Western blot. **(B)** MCF-7 cells were incubated with either control medium, or this medium containing Dp44mT (5 µM), DpC (5 µM), or 4-OHT (5 µM) for 24 h, followed by incubation for 15 min in the presence or absence of E_2_ (10 nM). Cellular lysates were then assessed for ER-α, p-ER-α^Ser167^, p-ER-α^Ser118^, PR, AR, and PRL-R *via* Western blot. **(C)** 3-dimensional (3D) spheroids composed of MCF-7 cells were grown for 48 h followed by 24 h treatment with control medium or this medium containing Dp44mT (5 µM) or DpC (5 µM) in the presence or absence of E_2_ (10 nM). Cellular lysates were then assessed for protein levels of ER-α, PR, AR, and PRL-R *via* Western blot. β-actin was used as a protein-loading control in (**A-C**). Results are mean ± SD (*n* = 3). Significant relative to untreated control cells: \**p* < 0.05, \*\**p* < 0.01, \*\*\**p* < 0.001. Significant relative to E_2_ untreated cells: ^#^*p* < 0.05, ^##^*p* < 0.01, ^###^*p* < 0.001.

The PR was detected as two isoforms, PR-A and PR-B [47], with both being quantified by densitometric analysis, as they have different, well-defined physiological functions and regulate different gene subsets [47]. Relative to the control, Dp44mT and DpC also significantly reduced the protein levels of the PR-A (90 kDa) and PR-B (118 kDa) isoforms in MCF-7 cells (**Fig. 2A**). Further, these agents also decreased AR and PRL-R expression (**Fig. 2A**), demonstrating for the first time their ability to inhibit multiple hormone receptors in BC. Incubation of MCF-7 cells with 4-OHT also markedly decreased PR-A, PR-B, AR, and PRL-R expression *versus* the control (**Fig. 2A**), in accordance with its potent anti-BC activity [48].

Further, the “gold standard,” clinically used iron chelator, DFO [49], was also able to significantly decrease ER-α, PR-A, AR, and PRL-R levels in MCF-7 cells (**Fig. 2A**), suggesting the well-characterized iron-binding activity of Dp44mT and DpC [17–20] was key to the activity observed herein. This conclusion was confirmed by the negative control thiosemicarbazone, Bp2mT, which cannot bind metal ions [17, 34–36] and had no significant effect on the expression of any of the proteins examined (**Fig. 2A**). To further validate this, we also probed for the well-established iron-regulated target, namely the metastasis suppressor, NDRG1, which is markedly up-regulated by these agents in an iron-dependent manner [23, 25]. In contrast to the pronounced down-regulation of ER-α, PR, AR, and PRL-R, NDRG1 was potently up-regulated following incubation with DFO, Dp44mT, or DpC in all cell-types examined (**Fig. 2A**), while the negative control, Bp2mT, had no significant effect (**Fig. 2A**). Assessing these proteins in T47D and ZR-75-1 cells (**Supplemental Fig. 1A, B**) revealed similar results as observed in MCF-7 cells, with Dp44mT and DpC significantly decreasing ER-α, PR, AR, and PRL-R protein levels relative to the control, being far more potent than 4-OHT.

As activation of ER-α is driven by E_2_ binding to this receptor followed by its phosphorylation at ER-α ^Ser167^ and ER-α^Ser118^ [13], studies next assessed the effect of preincubating BC cells for 24 h with Dp44mT (5 µM), DpC (5 µM), or 4-OHT (5 µM), and then subsequently adding E_2_ (10 nM) for 15 min/37°C (MCF-7) or 5 min/37°C (T47D). Examining the effect of Dp44mT and DpC on ER-α in MCF-7 cells in the presence or absence of E_2_, both agents caused a significant decrease in the expression of each ER-α isoform *versus* the control (**Fig. 2B**). Further, Dp44mT and DpC caused a pronounced and significant decrease in p-ER-α^Ser167^ levels in the absence of E_2_ relative to the control. While the addition of E_2_ increased phosphorylation at ER-α^Ser167^ and especially ER-α^Ser118^, this effect was completely inhibited by incubating cells with Dp44mT or DpC (**Fig. 2B**). Notably, 4-OHT also decreased p-ER-α^Ser167^ and p-ER-α^Ser118^ levels after adding E_2_ compared to the control, although to a far lesser extent than Dp44mT or DpC (**Fig. 2B**).

Dp44mT and DpC also significantly decreased PR-A (90 kDa) in the presence and absence of E_2_, having efficacy similar to 4-OHT (**Fig. 2B**). In the absence of E_2_, PR-B (118 kDa) was significantly decreased by Dp44mT, DpC, and 4-OHT *versus* the control, while in the presence of E_2_, only Dp44mT significantly decreased PR-B (118 kDa; **Fig. 2B**). Dp44mT and DpC also significantly decreased AR and PRL-R expression relative to the control in the absence and presence of E_2_ (**Fig. 2B**). The addition of E_2_ to the control significantly decreased PR-B, PR-A, AR, and PRL-R expression *versus* control cells without E_2_.

Assessing these alterations in protein expression and phosphorylation using T47D cells (**Supplemental Fig. 2A**) revealed similar results as observed in MCF-7 cells (**Fig. 2A, B**). In fact, Dp44mT and DpC significantly decreased ER-α, p-ER-α^Ser167^, PR-A, AR, and PRL-R protein levels in T47D cells in the presence and absence of E_2_ relative to the control, being generally more potent than 4-OHT (**Supplemental Fig. 2A**). Despite exhaustive attempts, p-ER-α^Ser118^ could not be detected in T47D cells even in the presence of E_2_ (**Supplemental Fig. 2A**). In contrast to the results observed with MCF-7 cells (**Fig. 2A, B**), 4-OHT did not significantly affect AR expression in T47D cells in absence of E_2_, while up-regulating PRL-R levels in the presence of E_2_ (**Supplemental Fig. 2A**).

To further confirm the effects of Dp44mT and DpC on these hormone receptors using a more pathophysiologically relevant model, MCF-7 cells were cultured as 3-dimensional spheroids and incubated with control medium or medium containing Dp44mT (5 µM) or DpC (5 µM) for 24 h in the absence or presence of E_2_ for the entire incubation (**Fig. 2C**). While Dp44mT and DpC significantly decreased ER-α, AR, and PRL-R expression *versus* the control in the presence and absence of E_2_, their effect on decreasing PR levels was lost when using 3D culture (**Fig. 2C**). To examine the effect of these agents on ER-α activation, the MCF-7 3D spheroids were incubated with Dp44mT or DpC for 24 h, followed by a shorter 15 min incubation with E_2_, as ER-phosphorylation occurs rapidly [16]. As also observed in 2D culture (**Fig. 2B**), Dp44mT and DpC significantly decreased total and phosphorylated ER-α levels in the presence or absence of E_2_ *versus* the control (**Supplemental Fig. 2B**).

Collectively, these studies demonstrate the significant decrease in expression of key BC drivers by Dp44mT and DpC in the presence and absence of E_2_ under 2D and 3D culture conditions in multiple BC cell-types.

### The proteasomal inhibitor, MG132, suppresses the ability of Dp44mT and DpC to down-regulate ER-α, PR, AR, and PRL-R protein levels

To determine the mechanism(s) by which thiosemicarbazones decrease ER-α, PR, AR, and PRL-R protein expression, initial studies assessed protein and mRNA levels of these receptors in MCF-7 cells following incubation with DpC from 0.25 h to 24 h (**Supplemental Fig. 3A, B**). As demonstrated in **Supplemental Fig. 3A**, ER-α protein levels were markedly and significantly decreased after a 6 h incubation with DpC *versus* the 0 h control time point. Similar to ER-α, the protein levels of PR, AR, and PRL-R were also significantly decreased after a 0.5 - 4 h incubation with DpC relative to the 0 h time point (**Supplemental Fig. 3A**).

**Figure 3.**
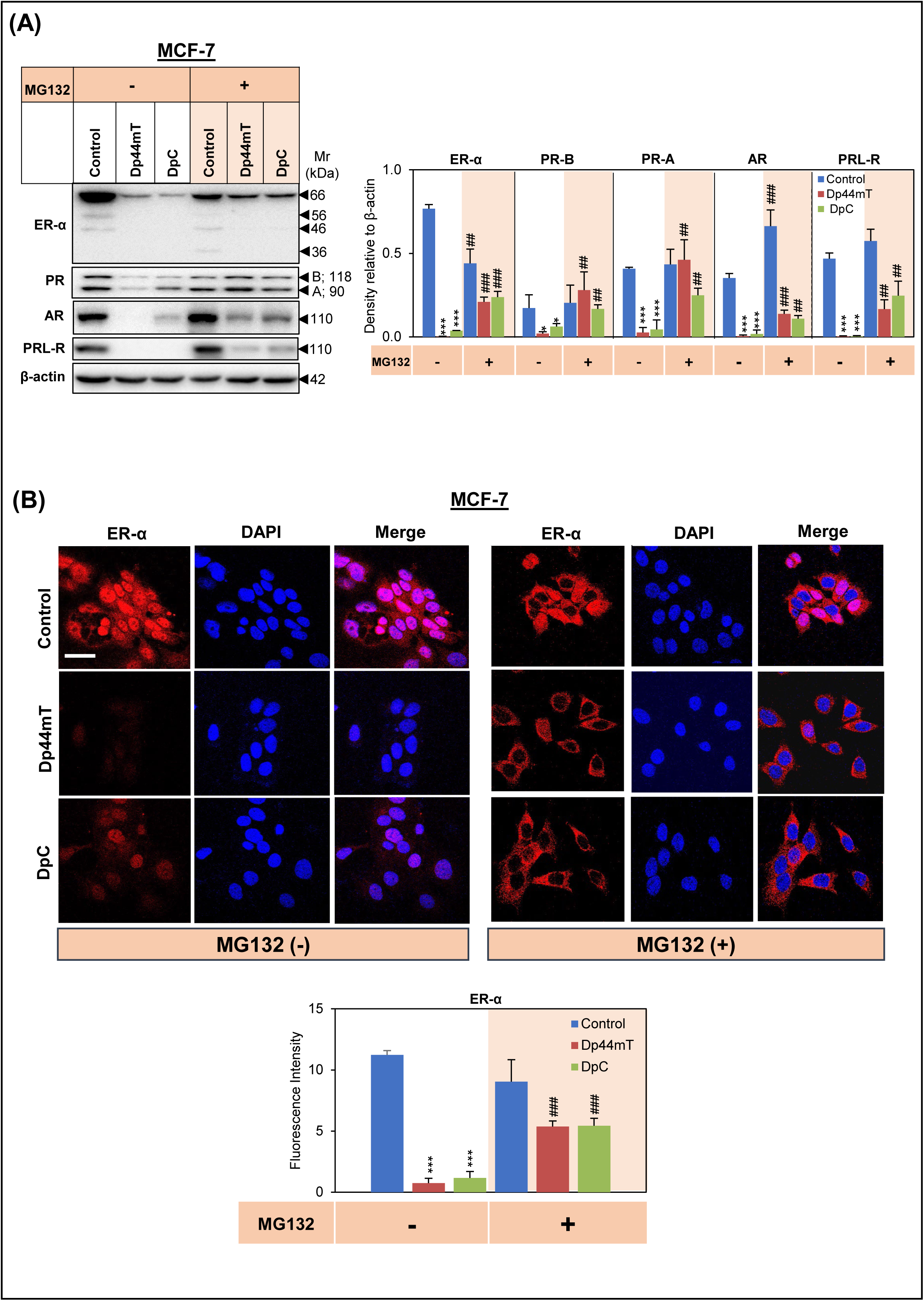
Dp44mT and DpC promote proteasomal degradation of ER-α, PR, AR, and PRL-R. **(A)** MCF-7 cells were incubated with either control medium or this medium containing DpC (5 µM) or Dp44mT (5 µM) in the presence or absence of MG132 (5 µM). Cellular lysates were then assessed for ER-α, PR, AR, and PRL-R protein levels *via* Western blot. β-actin was used as a protein-loading control. **(B)** MCF-7 cells were incubated with either control medium or this medium containing DpC (5 µM) or Dp44mT (5 µM) in the presence or absence of MG132 (5 µM) and assessed for ER-α expression *via* immunofluorescence microscopy. All images were taken using a 60x objective and were performed at the same exposure time using AxioVision™ software (Scale bar = 100 μm). Fluorescence intensity was quantified using ImageJ software. Results are mean ± SD (*n* = 3). Significant relative to untreated control cells: \**p* < 0.05, \*\*\**p* < 0.001. Significant relative to corresponding cells in absence of MG132: ^##^*p* < 0.01, ^###^*p* < 0.001.

Assessing the transcript levels of these receptors, *ER-α*, *AR,* and *PRL-R* expression was decreased after 8-24 h *versus* the control, while *PR-B* mRNA levels began to decline earlier at 4 h, recovering slightly before decreasing again at 24 h (**Supplemental Fig. 3B**). In contrast, *PR-A* mRNA expression did not significantly alter throughout the 24 h incubation. Overall, these results indicated that DpC decreased ER-α, PR, AR, and PRL-R protein levels to a much greater and more rapid extent prior to the decrease observed in their transcript levels, suggesting protein degradation was also involved in the activity of this agent.

Considering these latter studies, ER-α, PR, and AR proteins have been reported to be degraded predominantly *via* the ubiquitin-mediated proteasome pathway [50–52]. On the other hand, PRL-R degradation has been suggested to be mediated, at least in part, *via* the lysosome [31]. To investigate if Dp44mT and DpC promoted proteasomal degradation of these receptors, the well-characterized proteasomal inhibitor, MG132 [53], was utilized. In these studies, MCF-7 and T47D cells were incubated with Dp44mT (5 µM) or DpC (5 µM) alone or in combination with MG132 (at 2.5 or 5 µM for T47D or MCF-7 cells, respectively) for 24 h/37℃ and the protein levels of ER-α, PR, AR, and PRL-R examined. As previously demonstrated in **Figs. 1A, B** using MCF-7 cells, Dp44mT and DpC significantly decreased ER-α, PR, AR, and PRL-R levels relative to the control in the absence of MG132 (**Fig. 3A**). However, when combined with MG132, the effects of these agents on ER-α, AR, and PRL-R protein levels were significantly diminished. Moreover, the effects of Dp44mT and DpC on causing a pronounced and significant decrease in PR-A and PR-B protein levels were completely inhibited in the presence of MG132 (**Fig. 3A**).

Assessing MCF-7 cells *via* immunofluorescence microscopy revealed similar results as found for Western analysis, with ER-α levels being markedly and significantly reduced by Dp44mT and DpC in the absence of MG132, while this effect was significantly attenuated by adding MG132 (**Fig. 3B**). Notably, MG132 altered ER-α localization from predominantly nuclear to cytoplasmic, which has been reported to be due to proteasomal suppression that leads to cytosolic ER-α accumulation [54]. This latter observation in **Fig. 3B** acted as an appropriate positive control for the effective inhibition of proteasomal activity by MG132.

A similar effect to those observed using MCF-7 cells was also demonstrated for T47D cells, with the potent activity of Dp44mT and DpC at decreasing ER-α and AR protein expression being attenuated by MG132 (**Supplemental Fig. 4**). Interestingly, PR and PRL-R levels were markedly decreased by MG132 in control T47D cells and remained at these low levels in the presence of Dp44mT or DpC (**Supplemental Fig. 4**). Overall, these studies in **Figure 3** and **Supplemental Figures 3 and 4** suggest that together with the prominent decrease in mRNA levels, proteasomal function is also involved in the down-regulation of ER-α, PR, AR, and PRL-R by Dp44mT and DpC in MCF-7 cells.

**Figure 4.**
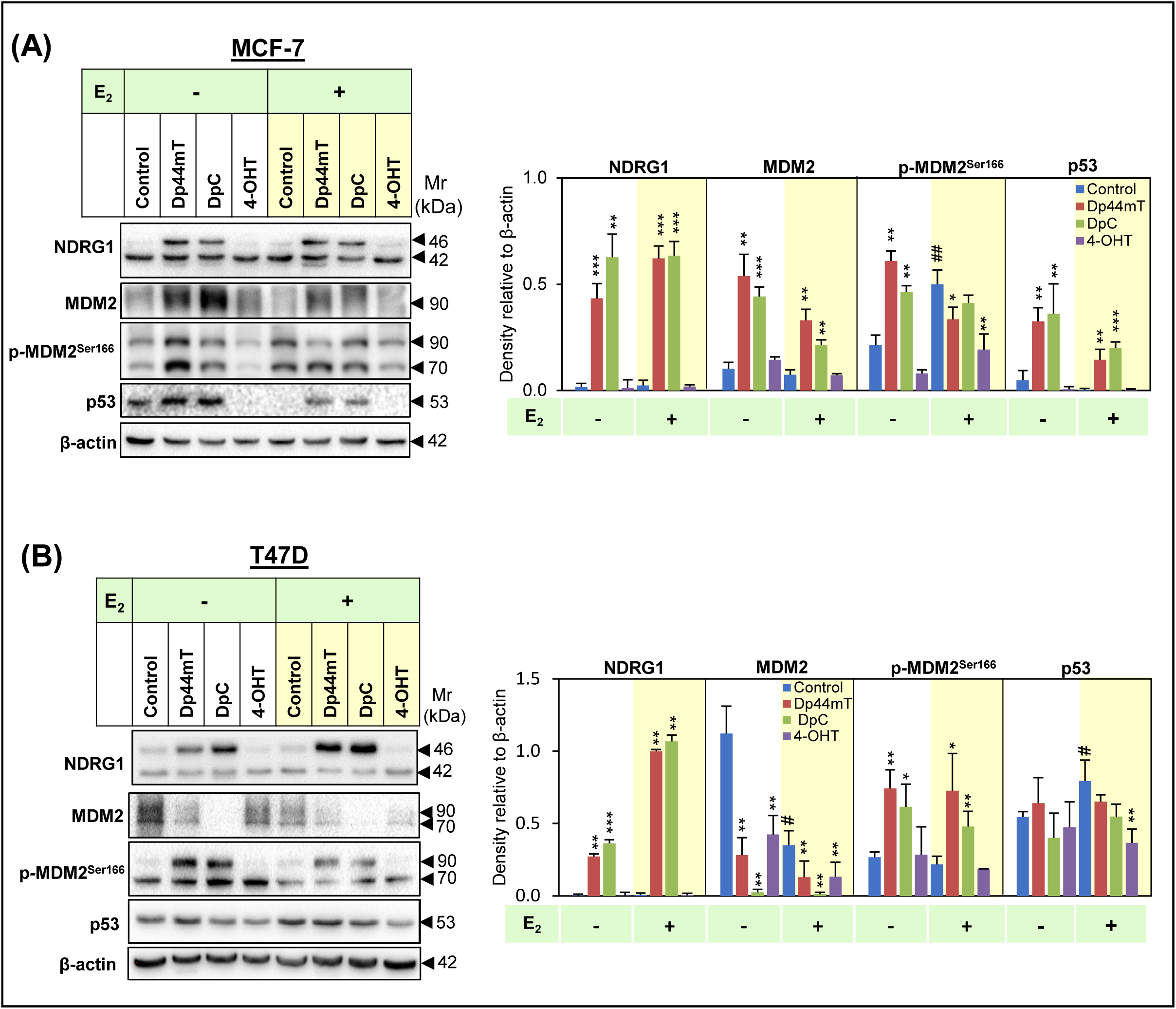
Effect of Dp44mT, DpC, and 4-OHT on NDRG1, MDM2, p-MDM2, and p53 protein levels in MCF-7 or T47D cells. **(A)** MCF-7 and **(B)** T47D cells were incubated with either control medium or this medium containing Dp44mT (5 µM), DpC (5 µM), or 4-OHT (5 µM) for 24 h in the presence or absence of E_2_ (10 nM). Cellular lysates were then assessed for the protein levels of NDRG1, MDM2, p-MDM2^Ser166^, or p53 *via* Western blot. β-actin was used as a protein-loading control. Results are mean ± SD (*n* = 3). Significant relative to untreated control cells: \**p* < 0.05, \*\**p* < 0.01, \*\*\**p* < 0.001. Significant relative to E_2_ untreated cells: ^#^*p* < 0.05, ^##^*p* < 0.01.

### Dp44mT and DpC up-regulate wild-type p53 and MDM2 in MCF-7 cells

Next, considering the results in **Figure 3** and **Supplemental Figure 4**, studies focused on proteins that promote proteasomal degradation of ER-α, including MDM2 and p53 [55]. Both MDM2 and p53 directly bind to ER-α and form a ternary complex that leads to ER-α ubiquitination and degradation [55]. In addition, MDM2 is a major regulator of functional p53 [56], and the phosphorylation at MDM2^Ser166^ is responsible for MDM2 translocation from the cytoplasm to the nucleus and, consequently, its activation [56]. Studies also assessed if the metastasis suppressor, NDRG1, that is potently induced by Dp44mT and DpC [17, 23] (**Fig. 2B**) could be involved in the ability of these agents to decrease ER-α expression. This hypothesis was important to investigate as NDRG1 is a critical target up-regulated by Dp44mT and DpC [18, 23, 24].

In **Figure 4A, B,** the expression of MDM2, p53, and NDRG1 was examined following a 24 h incubation with control medium or medium containing Dp44mT (5 µM), DpC (5 µM), or 4-OHT (5 µM) in the presence or absence of E_2_ (10 nM) in MCF-7 and T47D cells. Notably, MCF-7 cells express wild-type (WT) p53, while T47D cells have mutated p53 [57]. Both Dp44mT and DpC significantly increased NDRG1, MDM2, and p53 expression in MCF-7 cells *versus* the control in the presence or absence of E_2_, while 4-OHT had no significant effect (**Fig. 4A**). The ability of Dp44mT and DpC to up-regulate the tumor suppressor p53 is another marked therapeutic advantage. The up-regulation of WT-p53 is ascribed to the ardent ability of these thiosemicarbazones to bind cellular iron, as reported by our laboratory with other iron-binding drugs such as DFO [58, 59]. However, p-MDM2^Ser166^ levels were only significantly enhanced by Dp44mT and DpC in the absence of E_2_. Conversely, in the presence of E_2_, Dp44mT and 4-OHT significantly decreased p-MDM2^Ser166^ *versus* the control (**Fig. 4A**).

Examining the effects of Dp44mT and DpC using T47D cells revealed that while they induced a pronounced and significant increase in NDRG1 expression relative to the control, they potently decreased total MDM2 levels, while significantly increasing p-MDM2^Ser166^ *versus* the control in the presence and absence of E_2_ (**Fig. 4B**). Further, relative to the control, Dp44mT and DpC had no significant effect on p53 expression, probably because T47D cells have mutant p53 [57], which we demonstrated does not respond to cellular iron depletion [58, 59]. Incubation of T47D cells with 4-OHT also decreased total MDM2 expression in the presence and absence of E_2_ *versus* the control, while having no significant effect on p-MDM2^Ser166^ (**Fig. 4B**). These contrasting results between MCF-7 and T47D cells may be due to mutated p53 in T47D cells, which alters its effects on ER-α.

### Silencing *p53*, but not *NDRG1*, partly rescues the down-regulation of ER-α by Dp44mT

Since DpC and Dp44mT up-regulate NDRG1 expression in MCF-7 and T47D cells and WT-p53 in MCF-7 cells (**Fig. 4A, B**), it was hypothesized that these proteins may be involved in decreasing ER-α expression after incubation with these agents. While there are no studies showing how NDRG1 affects ER-α degradation, it is intriguing that ER-α levels were reported to be inversely correlated with NDRG1 in 96 patients [60]. Examination of p53 was also important, as it can: ***(i)*** directly bind to ER-α to alter its transcriptional activity; ***(ii)*** promote transcription of *ER-α*, leading to increased ER-α protein levels [61], and; ***(iii)*** form a complex with ER-α and MDM2 to promote ligand-dependent ER-α turnover [55]. To examine this hypothesis, studies implemented a negative control siRNA (si-Con) and a specific siRNA against *NDRG1* (si-NDRG1) in MCF-7 and T47D cells, followed by incubation with Dp44mT (5 µM) or DpC (5 µM) for 24 h/37°C (**Supplemental Fig. 5A, B**). Analogously, an siRNA against *p53* (si-p53) was utilized to silence *p53* in MCF-7 cells, followed by the same incubation conditions with Dp44mT or DpC (**Supplemental Fig. 6**).

**Figure 5.**
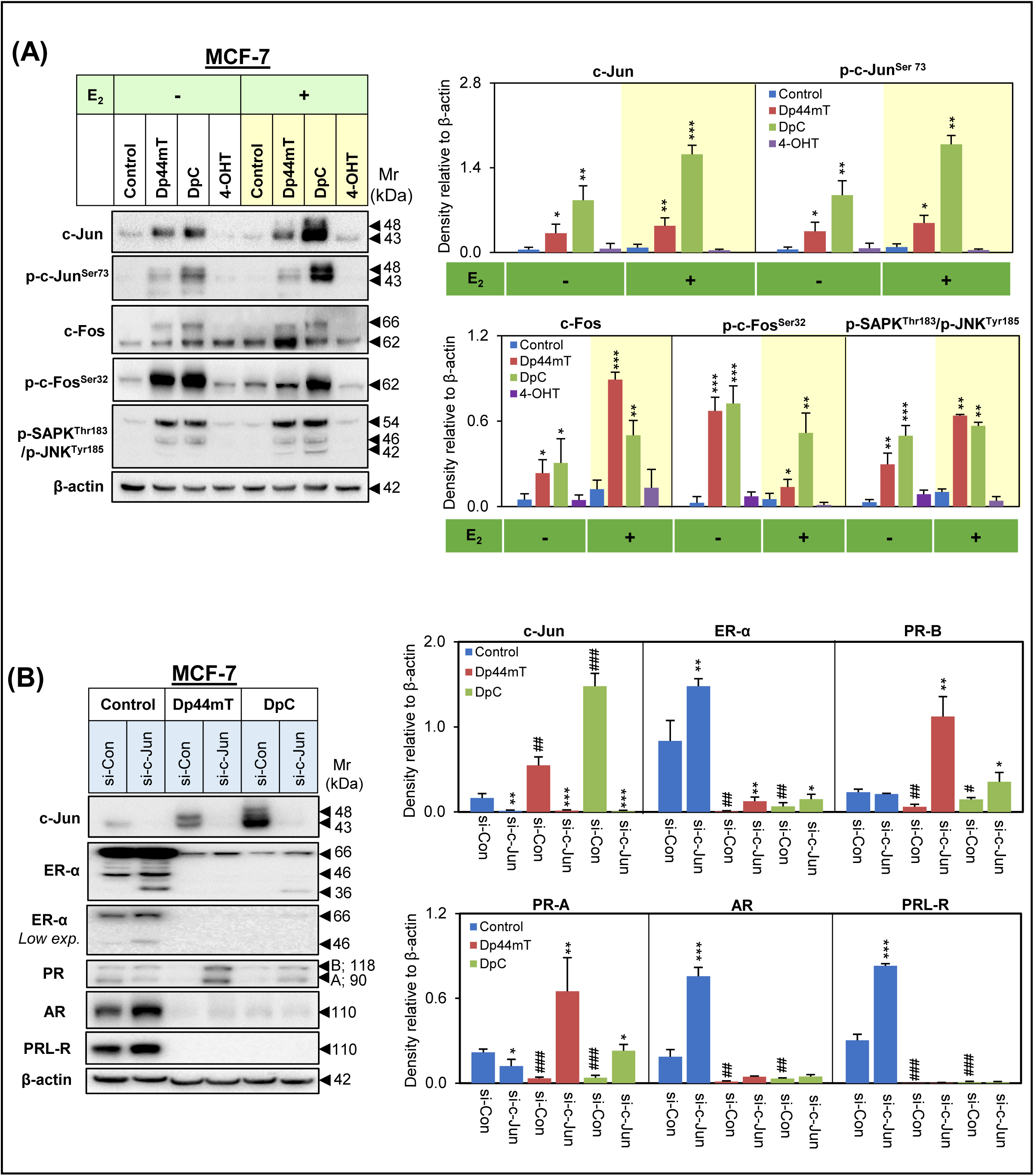
Dp44mT and DpC up-regulate c-Jun, c-Fos, and p-SAPK/p-JNK levels in MCF-7 cells. **(A)** MCF-7 cells were incubated with either control medium or this medium containing Dp44mT (5 µM), DpC (5 µM), or 4-OHT (5 µM) for 24 h in the presence or absence of E_2_ (10 nM). Cellular lysates were then examined for the protein levels of c-Jun, p-c-Jun^Ser73^, c-Fos, p-c-Fos^Ser32^, and p-SAPK ^Thr183^/p-JNK^Tyr185^ *via* Western blot. **(B)** MCF-7 cells were incubated with negative control siRNA (si-Con) or siRNA against c-Jun (si-c-Jun) for 24 h prior to treatment with Dp44mT (5 µM) or DpC (5 µM) for another 24 h. Cellular lysates were then assessed for protein levels of c-Jun, ER-α, PR, AR, and PRL-R *via* Western blot. β-actin was used as a protein-loading control in **(A)** and **(B)**. Results are mean ± SD (*n* = 3). Significant relative to si-Control cells: \**p* < 0.05, \*\**p* < 0.01, \*\*\**p* < 0.001. Significant relative to untreated control cells: ^#^*p* < 0.05, ^##^*p* < 0.01, ^###^*p* < 0.001.

**Figure 6.**
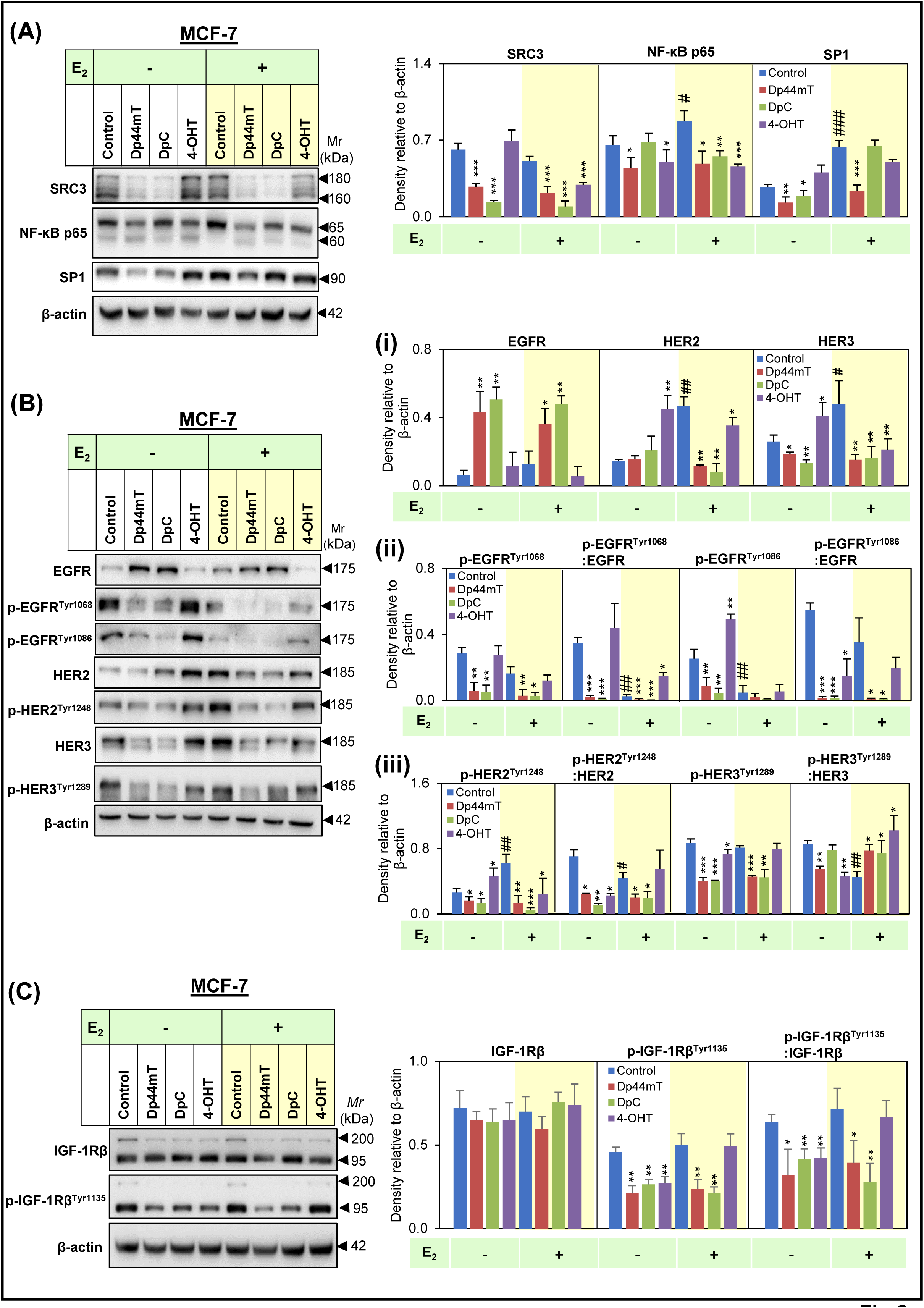
Dp44mT and DpC down-regulate co-activators, transcription factors and RTKs required for ER-α stability and signaling. MCF-7 cells were incubated with either control medium or this medium containing Dp44mT (5 µM), DpC (5 µM) or 4-OHT (5 µM) for 24 h in the presence or absence of E_2_ (10 nM). Cellular lysates were then assessed for protein levels of **(A)** SRC3, NF-κB p65, and SP1; **(B)** EGFR, p-EGFR^Tyr1068^, p-EGFR^Tyr1086^, HER2, p-HER2^Tyr1248^, HER3 and p-HER3^Tyr1289^; or **(C)** IGF-1Rβ and p-IGF-1Rβ^Tyr1135^. β-actin was used as a protein-loading control. Results are mean ± SD (*n* = 3). Significant relative to untreated control cells: \**p* < 0.05, \*\**p* < 0.01, \*\*\**p* < 0.001. Significant relative to cells not treated with E_2_: ^#^*p* < 0.05, ^##^*p* < 0.01, ^###^*p* < 0.001.

Silencing *NDRG1* under control conditions did not significantly affect ER-α expression, although there was a significant decrease in PR-A, PR-B, and AR expression in MCF-7 cells (**Supplemental Fig. 5A**) and a significant decrease of PR-A and AR in T47D cells relative to the si-Con (**Supplemental Fig. 5B**). However, silencing *NDRG1* in the presence of Dp44mT or DpC did not reverse the pronounced inhibitory effect of these agents on ER-α, PR, or AR expression in both cell-types (**Supplemental Fig. 5A, B**). These results suggest NDRG1 is not involved in the Dp44mT and DpC-mediated decrease in the expression of these proteins.

Silencing *p53* under control conditions using MCF-7 cells was accompanied by a significant reduction in ER-α, PR, AR, and PRL-R levels relative to the si-Con (**Supplemental Fig. 6**), demonstrating the ability of p53 to promote the expression of these proteins [61]. Moreover, silencing *p53* partially, but significantly rescued ER-α expression only in the presence of Dp44mT (**Supplemental Fig. 6**). This observation suggests that inhibition of p53 expression can partially reverse the activity of Dp44mT, but not DpC, in MCF-7 cells. Silencing *p53* in the presence of Dp44mT or DpC did not prevent their effects on inhibiting the expression of any of the other hormone receptors, suggesting their activity was independent of p53 (**Supplemental Fig. 6**).

In summary, these results in **Supplemental Figures 5** and **6** indicate that the up-regulation of NDRG1 by both thiosemicarbazones is not involved in the down-regulation of the receptors examined, while the effect of Dp44mT on decreasing ER-α protein expression is at least partially mediated by its ability to up-regulate p53.

### Dp44mT and DpC promote c-Jun and c-Fos activation

Another mechanism by which ER-α expression can be suppressed in cancer cells involves the transcription factor, c-Jun, which forms part of the AP-1 complex [62]. Recent studies have shown that c-Jun can directly suppress ER-α transcription by binding to the *ESR1* genomic locus [63]. Importantly, ER-α can also bind to the AP-1 complex at specific AP-1 response elements to regulate its transcriptional activity [62]. The downstream effects of the AP-1 transcription factor complex, which is composed of c-Jun and c-Fos DNA-binding proteins, can include either proliferation or apoptosis, depending on the cellular context [64]. Hence, the effects of Dp44mT (5 µM), DpC (5 µM), or OHT (5 µM) on c-Jun and c-Fos protein levels and their activating phosphorylations (p-c-Jun^Ser73^; p-c-Fos^Ser32^; [62]) were then examined in MCF-7 (**Fig. 5A**) and T47D cells (**Supplemental Fig. 7A**) after a 24 h incubation in the presence and absence of E_2_ (10 nM).

**Figure 7.**
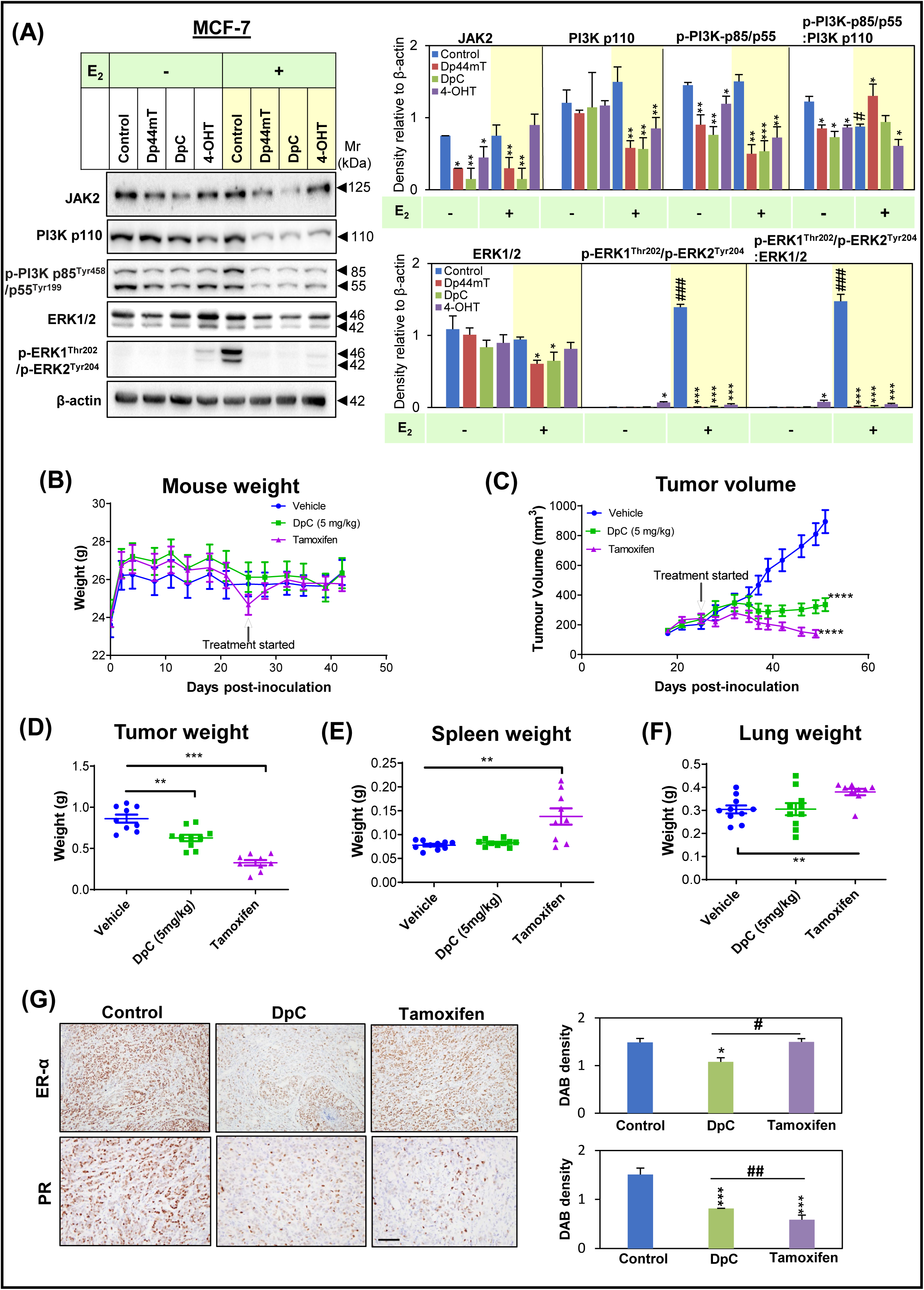
Dp44mT and DpC decrease downstream MAPK and PI3K signaling in MCF-7 cells and DpC inhibits BC growth *in vivo*. **(A)** MCF-7 cells were incubated with either control medium or this medium containing Dp44mT (5 µM), DpC (5 µM), or 4-OHT (5 µM) for 24 h in the presence or absence of E_2_ (10 nM). Cellular lysates were then assessed for levels of JAK2, PI3K p110, PI3K p85^Tyr458^/p55^Tyr199^, ERK1/2, and p-ERK1^Thr202^/p-ERK2^Tyr204^. β-actin was used as a protein-loading control. Results are mean ± SD (*n* = 3). Significant relative to the untreated control: \**p* < 0.05, \*\**p* < 0.01, \*\*\**p* < 0.001. Significant relative to cells not treated with E_2_: ^#^*p* < 0.05, ^###^*p* < 0.001. **(B-G)** Mice were orthotopically implanted in mammary fat pads with MCF-7 cells. Once palpable tumors of 200 mm^3^ formed, the animals were treated orally with either the vehicle control or DpC (5 mg/kg) 3x/week. tamoxifen slow-release pellets (35 mg, 60-day slow-release pellet) were implanted surgically for the tamoxifen-treated group (see *Materials and Methods*). The treatments were for 29 days and the mice were then sacrificed and examined for: (**B**) body weight; (**C**) tumor volume; (**D**) tumor weights; (**E**) spleen weights, and; (**F)** lung weights. Data presented in (**B**-**F**) are shown as the mean ± SD (*n* = 10 mice/group). (**G**) Immunohistochemistry of the tumor was performed to examine ER-α and PR protein expression. The scale bar represents 71 µm. Quantitation of staining was performed by ImageJ software. Results are shown as mean ± SD (*n* =5 images/slide; 10 tumor samples/group). Significance in comparison to vehicle control group: \**p* < 0.05, \*\**p* < 0.01, \*\*\**p* < 0.001, \*\*\*\**p* < 0.0001. Significance in comparison to DpC: #*p* < 0.05, ##*p* < 0.01.

As shown in **Fig. 5A**, Dp44mT and DpC significantly increased total and phosphorylated c-Jun and c-Fos in MCF-7 cells in the presence and absence of E_2_ relative to the control. Incubation with 4-OHT had no significant effect on the levels of c-Jun or c-Fos, or their phosphorylation in MCF-7 cells (**Fig. 5A**). Similar effects on c-Jun and p-c-Jun^Ser73^ were observed using T47D cells following treatment with Dp44mT and DpC (**Supplemental Fig. 7A**). However, only Dp44mT could significantly increase p-c-Fos^Ser32^ in T47D cells (**Supplemental Fig. 7A**). In contrast to MCF-7 cells, 4-OHT significantly increased or decreased c-Fos in the absence and presence of E_2_, respectively, relative to the respective control.

Considering that Dp44mT and DpC promoted activation of c-Jun and c-Fos, studies next examined p-SAPK^Thr183^/p-JNK^Tyr185^, which directly phosphorylates c-Jun and c-Fos in response to stress stimuli [65]. In fact, SAPK/JNK activation is critical for apoptosis in response to cellular stress [65]. As Dp44mT and DpC bind iron and copper resulting in redox-active complexes that induce cellular reactive oxygen species and stress [19, 21, 37], this may be a mechanism leading to c-Jun and c-Fos activation. In the presence and absence of E_2_, Dp44mT and DpC significantly increased p-SAPK^Thr183^/p-JNK^Tyr185^ in MCF-7 (**Fig. 5A**) and T47D cells (**Supplemental Fig. 7A**), while 4-OHT had no significant effect.

In summary, Dp44mT and DpC promote the AP-1 transcription factor complex *via* increased expression and activation of the c-Jun and c-Fos subunits.

### c-Jun is partly responsible for the effects of Dp44mT and DpC on down-regulating ER-α and PR

The increased c-Jun and p-c-Jun^Ser73^ levels in response to Dp44mT or DpC (**Fig. 5A; Supplemental Fig. 7A**) indicate that this protein may be involved in the ability of these agents to decrease *ER-α* mRNA levels (**Supplemental Fig. 3**) and subsequent protein expression after 24 h (**Fig. 2**; **Supplemental Figs. 1-2**). In fact, c-Jun was previously found to suppress ER-α expression by directly binding to the *ESR1* locus [63]. Further, *PR* and *AR* transcription have also been reported to be inhibited by the AP-1 complex [66], [67]. Hence, studies next examined whether the up-regulation of c-Jun observed with Dp44mT and DpC was responsible for their ability to decrease hormone receptor expression in BC.

To examine if c-Jun was involved in the Dp44mT- and DpC-mediated decrease of ER-α, PR, AR, and PRL-R expression, *c-Jun* was silenced in MCF-7 and T47D cells using a 48 h incubation with si-c-Jun, followed by incubation with Dp44mT or DpC (5 µM) for a further 24 h/37°C. As expected, c-Jun protein levels were markedly and significantly decreased in both cell-types transfected with si-c-Jun relative to the si-Con under all conditions (**Fig. 5B, Supplemental Fig. 7B**). Further, while Dp44mT and DpC significantly increased c-Jun expression in the si-Con-treated cells, this effect was potently inhibited in si-c-Jun-treated cells (**Fig. 5B, Supplemental Fig. 7B**). Under control conditions, *c-Jun* silencing was accompanied by significantly increased ER-α (see low exposure ER-α panel in **Fig.5B, Supplemental Fig. 7B**), AR, and PRL-R levels relative to the si-Con, demonstrating a role of c-Jun in promoting their suppression.

Assessing the effect of Dp44mT and DpC on ER-α expression, both agents caused a very pronounced decrease in ER-α levels in si-Con and si-c-Jun-treated cells *versus* the control (**Fig. 5B, Supplemental Fig. 7B**). However, this effect was slightly, yet significantly, less potent in the si-c-Jun-treated cells when compared to the si-Con for each condition examined (**Fig. 5B, Supplemental Fig. 7B**). This observation suggests that silencing *c-Jun* can slightly reverse the inhibitory effects of Dp44mT and DpC on ER-α expression.

The Dp44mT- and DpC-mediated decrease of PR expression in the si-Con was significantly reversed for PR-A and PR-B upon silencing *c-Jun* in MCF-7 cells (**Fig. 5B**) and partially reversed using T47D cells (**Supplemental Fig. 7B**). However, the inhibitory effects of Dp44mT and DpC on AR and PRL-R expression were not rescued when *c-Jun* was silenced in either cell-type (**Fig. 5B, Supplemental Fig. 7B**). These results indicated that the effects of Dp44mT and DpC on decreasing ER-α and PR expression are at least partially mediated by their ability to increase c-Jun expression. On the other hand, the effects of these agents on decreasing AR and PRL-R levels are independent of c-Jun.

Overall, the results from **Figs 3-5** and **Supplemental Figs 3-7** demonstrate that Dp44mT and DpC decrease the expression of ER-α, PR, AR, and PRL-R *via* the proteasomal degradation pathway and suppression of their mRNA levels. While the suppressive effects of these agents on ER-α and PR are partly mediated by p53 and c-Jun or c-Jun, respectively, the mechanisms involved in their pharmacological effects on AR and PRL-R remain elusive.

### Dp44mT and DpC down-regulate co-factors and transcription factors involved in ER-α signaling

To further mechanistically dissect how Dp44mT and DpC affect ER-α activation, stability, and signaling, studies then examined key proteins that function as: (**1**) co-activators of ER-α (*i.e.,* SRC3; [68]), or (**2**) transcription factors that interact with ER-α (*i.e.,* NF-κB p65, SP1; [69, 70]). Dp44mT and DpC significantly decreased the levels of SRC3, a key co-activator of ER-α, relative to the control in the presence or absence of E_2_ in MCF-7 cells (**Fig. 6A**). Similarly, NF-ᴋB p65 was significantly decreased by Dp44mT and 4-OHT *versus* the control in the absence of E_2_, while all agents significantly inhibited NF-ᴋB p65 expression in the presence of E_2_ (**Fig. 6A**). Expression of the transcription factor, SP1, was significantly decreased by Dp44mT in the presence and absence of E_2_, and by DpC in the absence of E_2_ only (**Fig. 6A**).

Similar results were also observed for T47D cells when examining SRC3 levels, which was significantly decreased by all agents compared to the control in the presence or absence of E_2_ (**Supplemental Fig. 8A**). However, NF-ᴋB p65 was significantly decreased by DpC and 4-OHT only in the presence of E_2_ in T47D cells. The expression of SP1 was also decreased by all agents in the absence of E_2_ and by Dp44mT and 4-OHT in the presence of E_2_ (**Supplemental Fig. 8A**). It was of interest that DpC up-regulated SP1 levels relative to the control in the presence of E_2_ in T47D cells.

**Figure 8.**
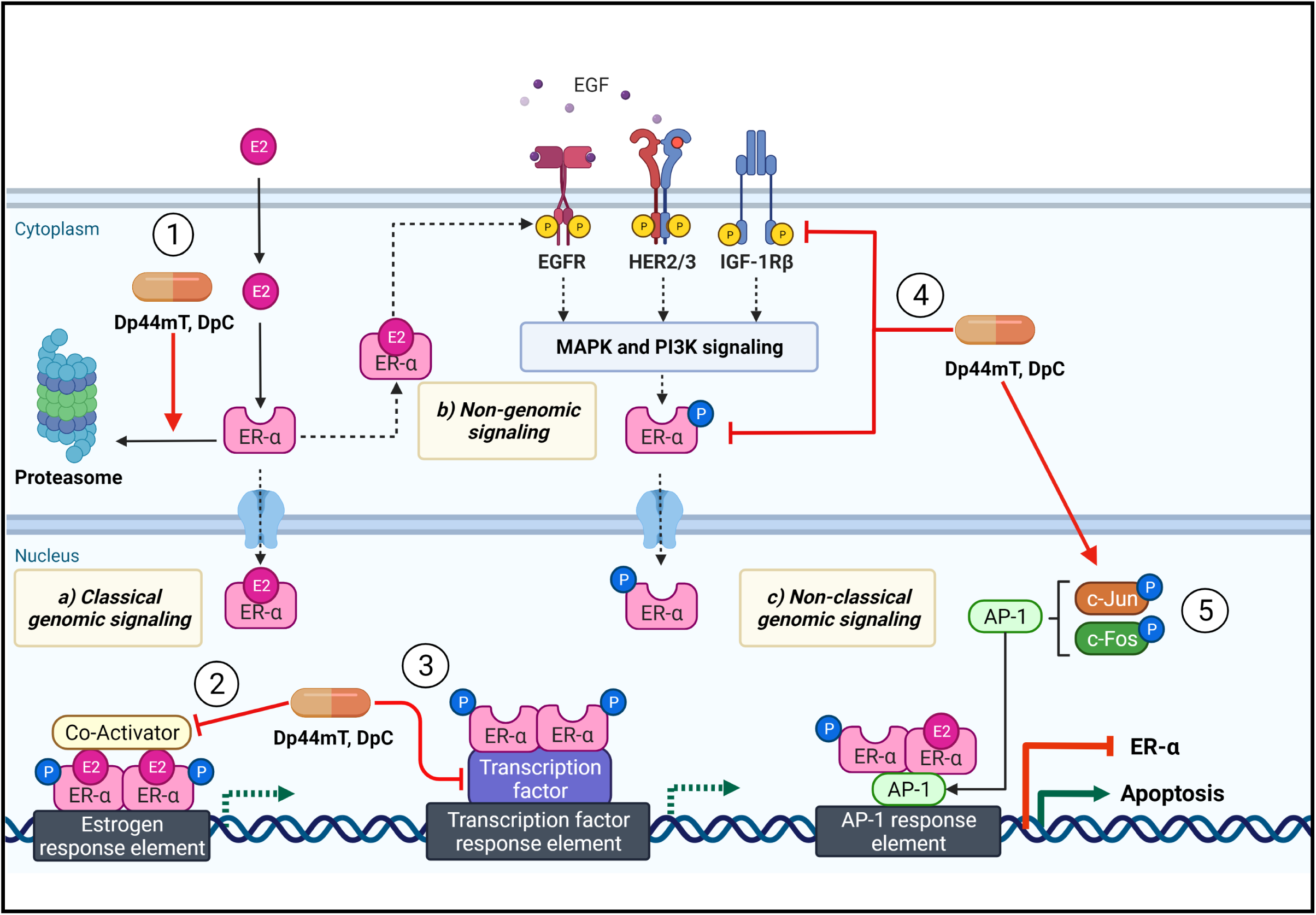
Schematic of the multi-faceted, comprehensive activity of DpC and Dp44mT on inhibiting ER-α signaling. Dp44mT and DpC were demonstrated to have multi-axial effects (**Effects 1-5**) on: **(a)** classical genomic signaling; **(b)** non-genomic signaling; and **(c)** non-classical genomic signaling. **(Effect 1)** Dp44mT and DpC inhibit classical genomic signaling by causing a pronounced decrease of ER-α protein levels and its phosphorylation at ER-α^Ser118^, indicating suppression of E_2_-dependent activation. This effect on decreasing ER-α protein levels included the ability of the thiosemicarbazones to promote proteasomal degradation of ER-α. **(Effect 2)** Dp44mT and DpC inhibit classical genomic ER-α signaling by inhibition of co-activators, such as SRC3. **(Effect 3)** Dp44mT and DpC inhibit classical genomic signaling by decreasing the expression of transcriptional factors (*i.e.,* NF-κB p65, SP1) that ER-α can bind to and promote oncogenic signaling. **(Effect 4)** Dp44mT and DpC inhibit non-genomic ER-α signaling by attenuating activation of RTKs such as EGFR, HER2/3, IGF-1Rβ, and downstream PI3K and MAPK signaling. **(Effect 5)** Dp44mT and DpC inhibit non-classical genomic signaling by promoting activation of c-Fos and c-Jun that form the AP-1 transcription factor complex that could induce apoptosis and decrease *ER-α* transcription.

Overall, these results in **Fig. 6A** and **Supplemental Fig. 8A** indicate that Dp44mT and DpC potently decrease SRC3 expression, and less consistently, NF-κB p65 and SP1 expression in MCF-7 and T47D cells, which could contribute to the observed decrease of ER-α expression after incubation with these agents.

### Dp44mT and DpC inhibit non-genomic ER-α signaling pathways *via* key RTKs

We next examined the effects of Dp44mT and DpC on major RTKs involved in crosstalk with ER-α to promote its activation and downstream signaling, including EGFR, HER2, HER3, and IGF-1Rβ [71, 72]. Once activated, these RTKs recruit JAK2 to their intracellular domains, directly facilitating activation of downstream PI3K/AKT and MAPK/ERK signaling cascades [73, 74]. Importantly, PI3K/AKT and MAPK/ERK signaling have been reported to directly promote ER-α phosphorylation and activation in the absence of E_2_ [75, 76].

While Dp44mT and DpC up-regulated total EGFR *versus* the control in the presence or absence of E_2_ in MCF-7 cells (**Fig. 6B(i)**), these agents markedly and significantly decreased EGFR phosphorylation at two activating sites, namely EGFR^Tyr1068^ and EGFR^Tyr1086^ (**Fig. 6B**). This was further evident by examining the ratios of p-EGFR to total EGFR, which were significantly decreased by Dp44mT and DpC relative to the control (**Fig. 6B(ii)**). Under control conditions, total HER2 expression was markedly and significantly up-regulated by E_2_ relative to the control without E_2_, with Dp44mT and DpC significantly decreasing HER2 in the presence of E_2_ *versus* the control (**Fig. 6B(i)**). In contrast, in the absence of E_2_, Dp44mT and DpC had no significant effect on HER2, while 4-OHT significantly increased HER2 (**Fig. 6B(i)**). Phosphorylated HER2^Tyr1248^, which indicates activation of this receptor [77], was significantly decreased by Dp44mT and DpC in the presence or absence of E_2_ *versus* the respective controls (**Fig. 6B(iii)**). Examining the ratio of p-HER2 ^Tyr1248^ to total HER2 also revealed a significant decrease with Dp44mT and DpC *versus* the control under all conditions (**Fig. 6B(iii)**). The expression of total HER3 (**Fig. 6B(i)**) and activated p-HER3^Tyr1289^ [78] (**Fig. 6B(iii)**) was also significantly decreased by Dp44mT and DpC compared to the respective controls in the presence or absence of E_2_. Assessing the ratio of p-HER3 to total HER3 revealed that Dp44mT, DpC, and 4-OHT up-regulated this ratio relative to the respective control in the presence of E_2_ (**Fig. 6B(iii)**). These results suggested that the inhibitory effect of these agents is on the total levels of this protein rather than its activation.

Examining the EGFR family of receptors in T47D cells, similar results to those found using MCF-7 cells (**Fig. 6B**) were observed (**Supplemental Fig. 8B**). In fact, Dp44mT and DpC significantly decreased p-EGFR^Tyr1068^, p-EGFR^Tyr1068^: EGFR ratio, p-EGFR^Tyr1086^, p-EGFR^Tyr1086^: EGFR ratio, HER2, p-HER2^Tyr1248^, p-HER2^Tyr1248^:HER2 ratio, and p-HER3^Tyr1289^ *versus* the control in the presence or absence of E_2_, being generally more potent than 4-OHT (**Supplemental Fig. 8B(i-iii)**). In the absence of E_2_, 4-OHT significantly increased EGFR phosphorylation at Tyr1068 and Tyr1086 relative to the control (**Supplemental Fig. 8B(ii)**). Both Dp44mT and DpC also significantly decreased total HER3 relative to the control in the presence of E_2_ (**Supplemental Fig. 8B(i)**). Overall, these results demonstrate the potent inhibitory effects of Dp44mT and DpC on activation of the EGFR family of receptors.

Relative to the control, total IGF-1Rβ expression (detected as both the pre-IGF-1R (200 kDa) and IGF-1Rβ (95 kDa) bands) was not significantly affected by any of the agents examined in MCF-7 cells (**Fig. 6C**). However, the activating phosphorylation at IGF-1Rβ^Tyr1135^ was significantly decreased by Dp44mT and DpC in the presence or absence of E_2_ (**Fig. 6C**). Similarly, the ratio of p-IGF-1Rβ to total IGF-1Rβ also showed a significant decrease after incubation with Dp44mT and DpC relative to the control (**Fig. 6C**), suggesting these agents inhibit activation of this protein rather than its total levels in MCF-7 cells. Examining T47D cells (**Supplemental Fig. 8C**), both total IGF-1Rβ and p-IGF-1Rβ^Tyr1135^ were significantly decreased by Dp44mT and DpC in the presence and absence of E_2_ relative to the control. This effect resulted in no significant alteration in the p-IGF-1Rβ^Tyr1135^: IGF-1Rβ ratio *versus* the control in the absence of E_2_, or a significant increase in the presence of E_2_, respectively.

Examining downstream signaling pathways in MCF-7 cells activated by EGFR, HER2, HER3, and IGF-1Rβ, it was demonstrated that Dp44mT and DpC resulted in a significant decrease in JAK2 expression *versus* the control in the presence or absence of E_2_ (**Fig. 7A**). In contrast, 4-OHT was less effective, and slightly, but significantly, decreased JAK2 relative to the control in the absence of E_2_, while having no significant activity in its presence.

JAK2 is a non-receptor protein tyrosine kinase that activates multiple downstream pathways, including PI3K/AKT and MAPK/ERK to promote the survival and migration of ER-α positive BC cells [73, 74]. In the presence of E_2_, Dp44mT and DpC significantly decreased PI3K p110 and p-PI3K p85^Tyr458^/p55^Tyr199^ *versus* the control (**Fig. 7A**). Considering this, in the presence of E_2_, the ratio between p-PI3K p85^Tyr458^/p55^Tyr199^ and PI3K p110 was significantly increased or not altered by Dp44mT or DpC, respectively. On the other hand, in the absence of E_2_, none of the agents had any significant effect on total PI3K p110 levels, but they all significantly decreased p-PI3K p85^Tyr458^/p55^Tyr199^ relative to the control (**Fig. 7A**). Under these latter conditions, in the absence of E_2_, the ratio between p-PI3K p85^Tyr458^/p55^Tyr199^ and PI3K p110 was significantly decreased by all agents *versus* the control, indicating a decrease in phosphorylation rather than total protein levels (**Fig. 7A**).

Examining total ERK1/2 expression, Dp44mT and DpC significantly decreased their levels in the presence of E_2_ *versus* the control, while having no effect in the absence of E_2_ (**Fig. 7A**). The phosphorylation of ERK1^Thr202^/ERK2^Tyr204^, which enables their activation [79], was barely evident in the absence of E_2_ under all conditions, but was potently increased in control cells by E_2_ (**Fig. 7A**). However, this marked activation of ERK1/2 by E_2_ was almost completely abolished by Dp44mT, DpC, or 4-OHT, resulting in a marked and significant decrease in the p-ERK1^Thr202^/p-ERK2^Tyr204^ to total ERK1/2 ratio relative to the control (**Fig. 7A**).

Overall, Dp44mT and DpC decreased the expression and/or activation of multiple oncogenic proteins in BC cells that interact with and stimulate ER-α activity, particularly in the presence of E_2_.

### DpC is well tolerated *in vivo*, suppresses MCF-7 orthotopic xenograft growth, and decreases ER-α and PR expression in the tumor

Considering DpC has far greater tolerability than Dp44mT *in vivo* [18, 21, 22], we further assessed the efficacy of DpC as a new therapeutic strategy against BC using MCF-7 orthotopic xenografts (**Fig. 7B-G**). As MCF-7 cells require E_2_ to proliferate and form tumors *in vivo*, these mice were also implanted with slow-release E_2_ pellets to sustain tumor growth, which was done 4 days before inoculation with cancer cells [80].

Once the tumors were palpable (23 days after implantation of MCF-7 cells), mice were treated with either vehicle control, DpC (5 mg/kg), or a tamoxifen slow-release pellet (35 mg pellets with 60-day release [80]). After the tamoxifen pellet was implanted, a transient but significant total body weight reduction was observed (see “treatment started” arrow in **Fig. 7B**), while DpC was well tolerated and led to no weight loss (**Fig. 7B**). After 51 days, the tumor size in mice treated with the vehicle control reached an average volume of 975 ± 118 mm^3^, while DpC and tamoxifen-treated groups had significantly reduced tumor volumes to 359 ± 60 mm^3^ and 152 ± 3 mm^3^, respectively (**Fig. 7C**). The reduction in tumor volume became apparent after only 12 days of treatment with both agents. Examining the tumor weight at the end of the study again demonstrated a pronounced and significant reduction in tumor weight by DpC and tamoxifen (627 ± 124 mg and 373 ± 176 mg, respectively) *versus* the vehicle control (793 ± 260 mg; **Fig. 7D**).

To assess the tolerability of both treatments, organs were harvested from mice after euthanasia, and spleen and lung weights were measured. We observed no significant difference in spleen and lung weight when treated with DpC compared to the vehicle control (**Fig. 7E, F**). Additionally, there were no signs of breathing or pulmonary distress in the DpC-treated mice, while tamoxifen significantly increased spleen and lung weight (**Fig. 7E, F**) and led to breathing distress. Hence, tamoxifen showed more potent anti-tumor efficacy, while also being less well tolerated than DpC. On the other hand, DpC demonstrated high tolerability, suggesting its potential as a novel treatment strategy for ER-positive BC.

The tumors from control and treated mice were further assessed for ER-α and PR expression *via* immunohistochemistry, where DpC significantly reduced the expression of ER-α and PR *in vivo* (**Fig 7G**). The expression of ER-α was also significantly decreased by DpC compared to tamoxifen, with tamoxifen having no significant effect on ER-α expression. Both DpC and tamoxifen significantly decreased PR expression *versus* the control, with tamoxifen demonstrating slightly, but significantly greater activity than DpC. The expression of AR was not detected in these tumor specimens, despite exhaustive attempts and multiple conditions being trialed.

## Discussion

There is an urgent need for innovative therapeutics with new mechanisms of action to counter the increasing incidence of resistance to BC endocrine therapy and the highly aggressive disease associated with it [11]. The significance of the current investigation is underlined by the fact that the most common BC-type, ER-positive BC, is driven by ER-α and is also characterized by PR, AR, PRL-R, and tyrosine kinase expression, which act integrally with ER-α to promote deadly BC progression [5–9]. DpC and Dp44mT disrupt these multiple, key inter-receptor interactions through a novel mechanism targeting redox-active metals, decreasing the expression and/or activation of these proteins rather than inhibiting ER-α alone (as with tamoxifen). Hence, these agents constitute an innovative pharmacological strategy for BC treatment.

The critical, well-established primary molecular targets of DpC and Dp44mT are tumor cell iron and copper [17–20]. In agreement with this, the suppressive effects of these agents on hormone receptor protein levels were totally ablated by inactivation of the metal-binding site, as demonstrated using the especially designed negative control thiosemicarbazone, Bp2mT [17, 34–36]. This conclusion was substantiated by examining the clinically used gold standard iron chelator, DFO, which led to similar down-regulation of these key hormone receptors as DpC and Dp44mT. Other key anti-oncogenic benefits of targeting tumor cell iron pools were the marked induction of the metastasis suppressor, NDRG1 [23–25], which inhibits deadly BC metastasis *in vivo* [26], and the tumor suppressor, WT-p53, both of which are regulated by cellular iron levels [23-25, 58, 59]. Significantly, the up-regulation of p53 was linked to an apoptotic gene signature (**Fig. 1B**) that is known to lead to tumor cell death with these thiosemicarbazones [18, 19, 21]. Indeed, after complexation with iron and copper, Dp44mT and DpC become redox-active and cytotoxic, being selectively more active against cancer cells *versus* their normal counterparts [20, 22].

As part of their unique multi-axial mechanism of action, Dp44mT and DpC suppressed three arms of ER-α signaling, having multiple effects on **(a)** classical estrogen-dependent genomic signaling; **(b)** non-genomic signaling; and **(c)** non-classical genomic signaling (**Fig. 8, Effects 1-5**). First, considering the effect on classical genomic signaling, Dp44mT and DpC caused a pronounced decrease in ER-α protein levels (**Fig. 8, Effect 1**) and p-ER-α^Ser118^, indicating suppression of E_2_-dependent activation [81]. Additionally, these agents inhibited the expression of co-activators (*i.e.,* SRC3; (**Fig. 8, Effect 2**) and transcription factors (*i.e.,* SP1, NF-κB p65) directly associated with ER-α transcription [68–70] (**Fig. 8, Effect 3**). Second, regarding non-genomic signaling, Dp44mT and DpC inhibited EGFR, HER2, HER3, and IGF-1Rβ activation and their downstream MAPK and PI3K pathways (**Fig. 8, Effect 4**), which can be rapidly activated by E_2_ *via* ER-α [71, 72]. Dp44mT and DpC also decreased p-ER-α^Ser167^ levels, a known downstream target of MAPK and PI3K signaling [75, 76]. Third, in terms of non-classical genomic signaling (**Fig. 8, Effect 5**), Dp44mT and DpC promoted activation of c-Jun and c-Fos that form the AP-1 transcription factor complex, which could induce apoptosis and reduce *ER-α* transcription [62]. Collectively, DpC and Dp44mT resulted in comprehensive, multi-faceted inhibition of ER-α signaling.

Considering further non-classical genomic signaling of ER-α (**Fig. 8, Effect 5**), Dp44mT and DpC also increased p-SAPK^Thr183^/p-JNK^Tyr185^ levels that are responsible for AP-1 complex activation [64]. Upon cellular stress, such as that induced by reactive oxygen species, the AP-1 complex promotes apoptosis [62]. Hence, activation of AP-1 in response to Dp44mT and DpC, which are highly redox-active after complexation with iron or copper [19, 21, 37], will likely lead to the apoptosis observed with these agents [18, 19, 21]. This proposition is further supported by our RNAseq data indicating increased expression of apoptosis-related genes by DpC in MCF-7 cells (*i.e., p53*, *GADD45*, *BAX*, *PUMA*, *p21, etc*.; **Fig 1B**).

In fact, we previously demonstrated that Dp44mT could enhance JNK and p38 MAPK signaling pathways [82], which promotes AP-1 mediated apoptosis [83].

The AP-1 complex inhibits *ER-α* mRNA transcription [63], which could contribute to the decreased *ER-α* mRNA levels, and thus, protein expression observed in response to Dp44mT and DpC. As ER-α itself can regulate transcription of *PR*, *AR,* and *PRL-R* [84–87], and its own transcription *via* AP-1 [63], the reduced ER-α protein levels are likely a significant contributor to the subsequently decreased expression of these hormone receptors in response to these thiosemicarbazones. Further, these latter suppressive effects on *ER-*α mRNA expression were also supplemented by the ability of Dp44mT and DpC to inhibit the expression of co-activators and transcription factors that promote ER-α transcriptional activity, including SRC3, NF-κB p65, and SP1. Apart from the decrease in mRNA levels, another mechanism by which Dp44mT and DpC inhibit ER-α, PR, AR, and PRL-R expression involved proteasomal degradation. Our RNAseq analysis revealed that DpC could modulate a plethora of proteins involved in the positive regulation of proteasomal protein catabolism in MCF-7 cells (*i.e.*, *HERPUD1*, *RNF19B*, *etc.*; **Supplemental Table 2**), providing important clues for future studies. Additionally, the decrease in ER-α and PR protein expression by the thiosemicarbazones was also partially mediated by the up-regulation of p53 and c-Jun or c-Jun, respectively. Together, a complex interplay of regulatory mechanisms at the mRNA and protein levels was responsible for the down-regulation of the hormone receptors by Dp44mT and DpC.

Non-genomic and non-classical E_2_-independent genomic ER-α signaling are major gateways towards tamoxifen resistance in ER-positive BC [88, 89]. In fact, activation of EGFR, HER2, HER3, and IGF-1Rβ are vital for multiple downstream oncogenic pathways crucial for driving BC progression, metastasis, and endocrine resistance [90–94]. Hence, the ability of Dp44mT and DpC to inhibit these additional arms of ER-α signaling may have important implications for overcoming tamoxifen resistance. Considering this, we reported that combining DpC or Dp44mT with tamoxifen and various other chemotherapeutics demonstrates synergistic anti-proliferative activity against BC cells [95–97]. This beneficial activity could assist in overcoming tamoxifen resistance, which is a major clinical hurdle in treating deadly ER-positive BC [11]. In fact, our previous studies *in vitro* demonstrated that DpC could overcome tamoxifen resistance in BC cells [95, 96]. While hormonal therapies such as tamoxifen are well-established for treating ER-positive BC, the results herein demonstrate for the first time the surprising activity of these agents mediated by binding metals on the expression/activation of multiple key receptors integrally involved in facilitating BC pathogenesis [5–9].

The current investigation demonstrates the efficacy and safety of DpC in inhibiting BC growth *in vivo*, using an orthotopic xenograft of MCF-7 cells. DpC was chosen for the *in vivo* study as it demonstrated superior tolerability than Dp44mT [22], was trialed clinically as an anti-cancer drug [32], and has never been assessed against BC tumors *in vivo*. DpC potently inhibited BC tumor growth *in vivo* with very good tolerability, resulting in decreased ER-α and PR protein expression in BC tumors. Although DpC was not as potent as tamoxifen in reducing tumor growth, it was better tolerated than tamoxifen. Additionally, considering the ability of DpC to inhibit multiple key pathways [95–98] that promote endocrine resistance also suggests it could be a superior strategy for BC treatment that potentially overcomes tamoxifen resistance [95]. Traditional drug design in terms of cancer therapy dictates 1 drug with 1 target that unfortunately leads to deadly resistance, such as with tamoxifen [32]. This investigation is unique in terms of the concept of innovative and safe, multi-targeted drugs that attack oncogenic interactions between multiple key receptors known to play critical roles in the progression of ER-positive BC [5–9].

In summary, the current research demonstrates an innovative, non-hormonal pharmacological strategy using redox-active Dp44mT and DpC to markedly inhibit ER-α, PR, AR, PRL-R, and protein tyrosine kinase expression/activation (including JAK2, EGFR, HER2, HER3, and IGF-1Rβ) in luminal A and luminal B BC cell-types. The activation of ER-α in response to E_2_ was also inhibited by these agents resulting in suppression of critical downstream signaling. These agents decreased the expression and/or activation of multiple co-factors and transcription factors, namely SRC3, SP1, and NF-κB p65, that promote ER-α activity, and DpC was effective and highly tolerable *in vivo* against the growth of ER-α positive tumors. This study highlights that through a novel, non-hormonal mechanism targeting redox active metal ions, Dp44mT and DpC disrupt multiple fundamental inter-receptor interactions between PR, AR, PRL-R, and tyrosine kinases that act integrally with ER-α to promote BC. As such, these agents constitute an innovative therapeutic approach.

## Statements and Declarations

### Author contributions

All authors contributed to the study’s conception and design. Material preparation, data collection, and analysis were performed by Faten Shehadeh-Tout, Heloisa H. Milioli, Suraya Roslan, Mahendiran Dharmasivam, Tharushi Wijesinghe and Mahan Gholam Azad. The first preliminary draft of the manuscript was written by Faten Shehadeh-Tout and all authors commented on previous versions, with extensive revisions by Des R. Richardson and Zaklina Kovacevic. All authors read and approved the final manuscript. Zaklina Kovacevic and Des R. Richardson provided all financial support for the investigation through grant support as co-chief investigators and supervisors.

### Data availability

The datasets generated during and/or analysed during the current study are available in the GEO repository (GEO accession number GSE192942).

### Ethics approval

This study was performed in line with the principles of the Declaration of Helsinki. Approval was granted by the Ethics Committee of the Olivia Newton-John Cancer Research Institute, Melbourne, Victoria.

