## Supplemental Figures for "An Innovative Non-Hormonal Strategy Targeting Redox Active Metals to Down-Regulate Estrogen-, Progesterone-, Androgen- and Prolactin-Receptors in Breast Cancer"

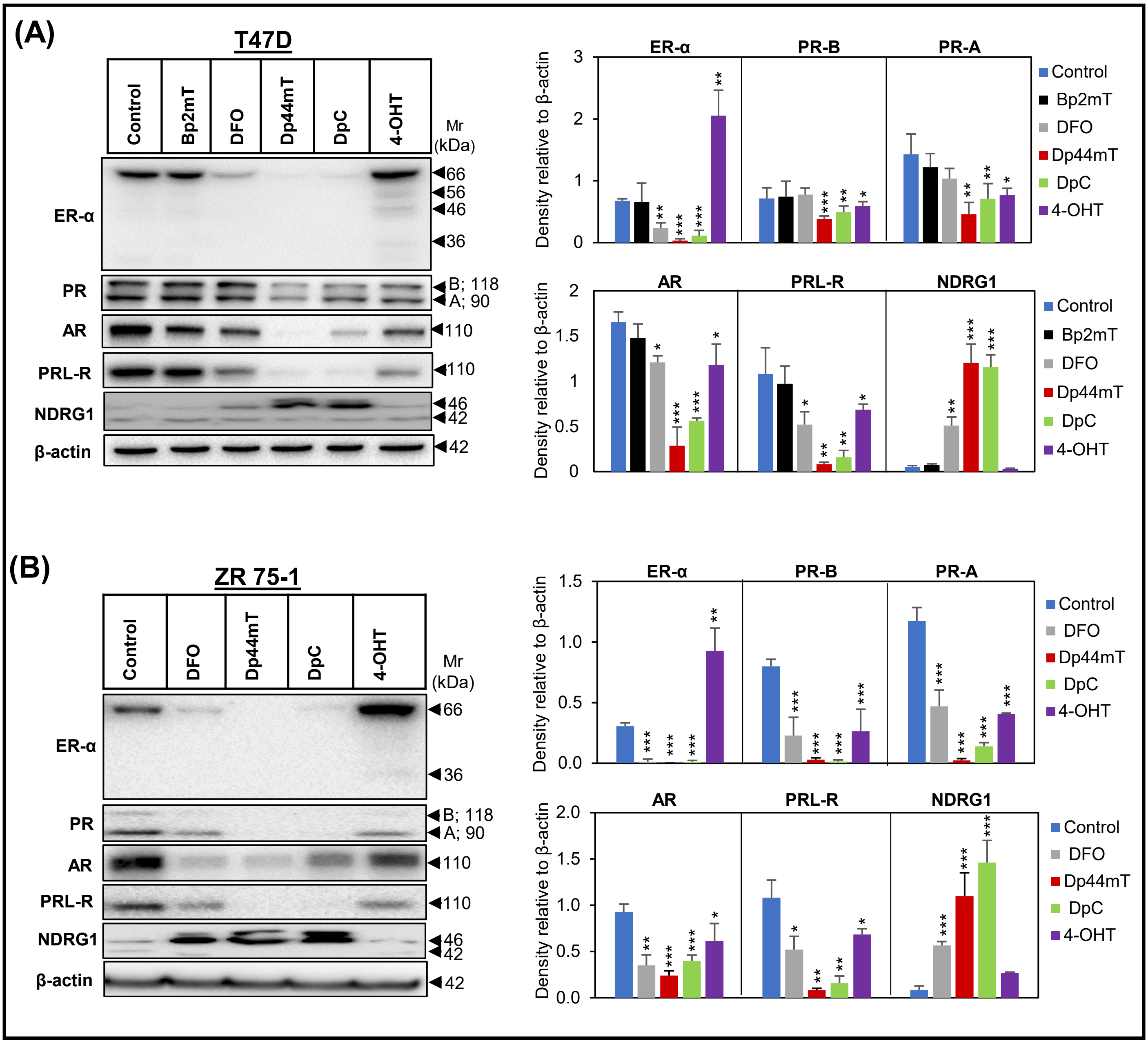


**Supplemental Figure 1**

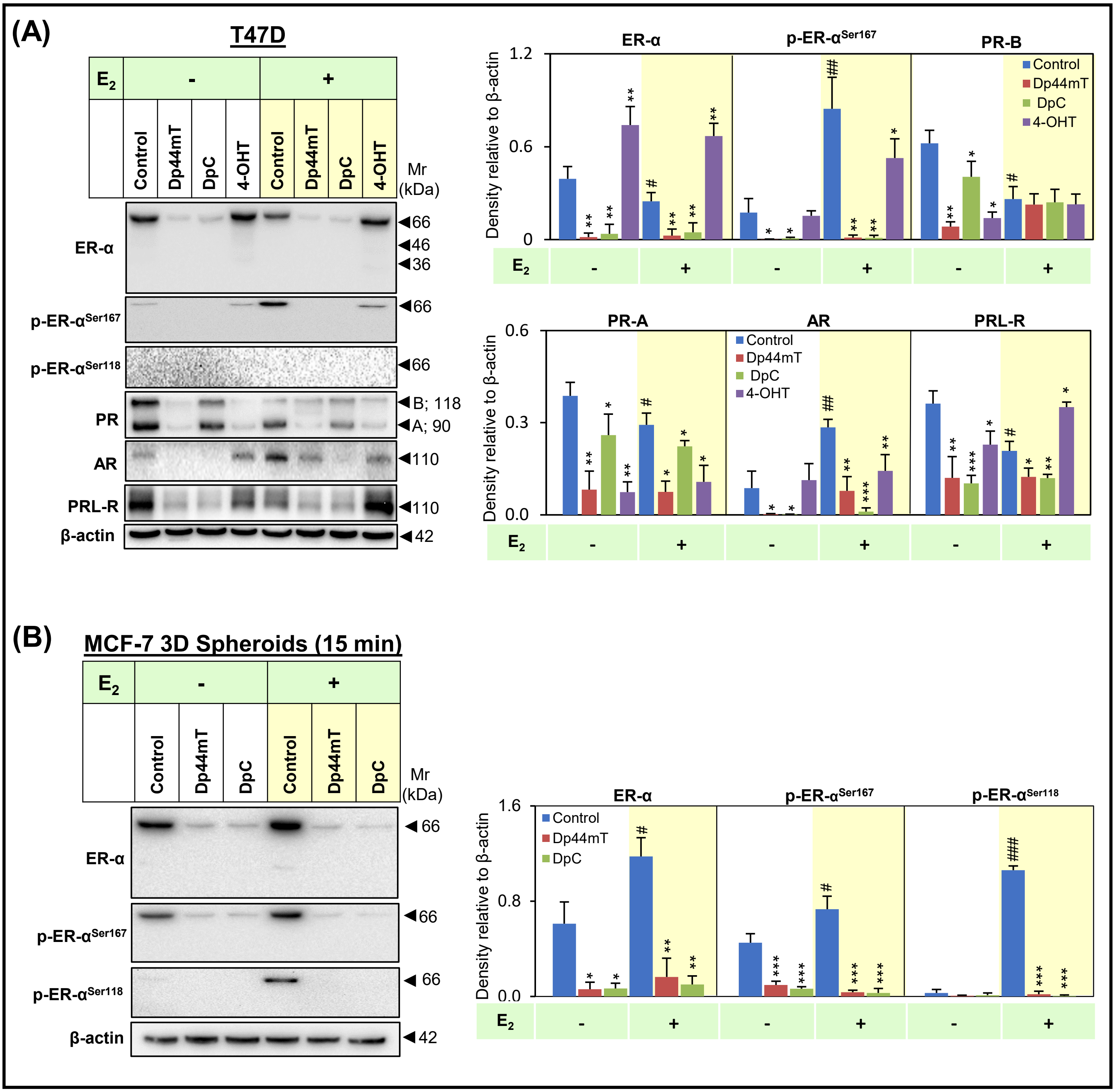


**Supplemental Figure 2**

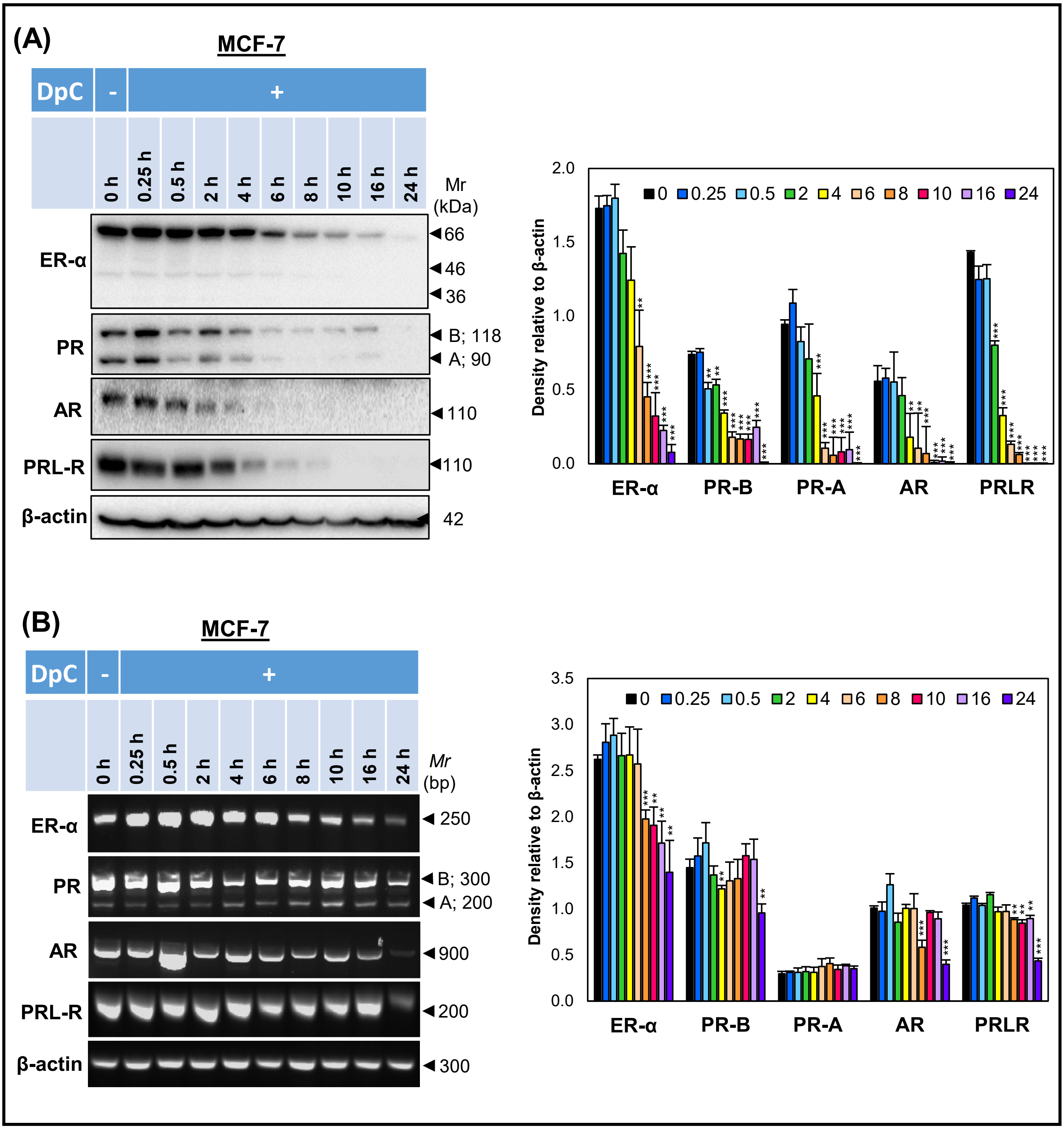


**Supplemental Figure 3**

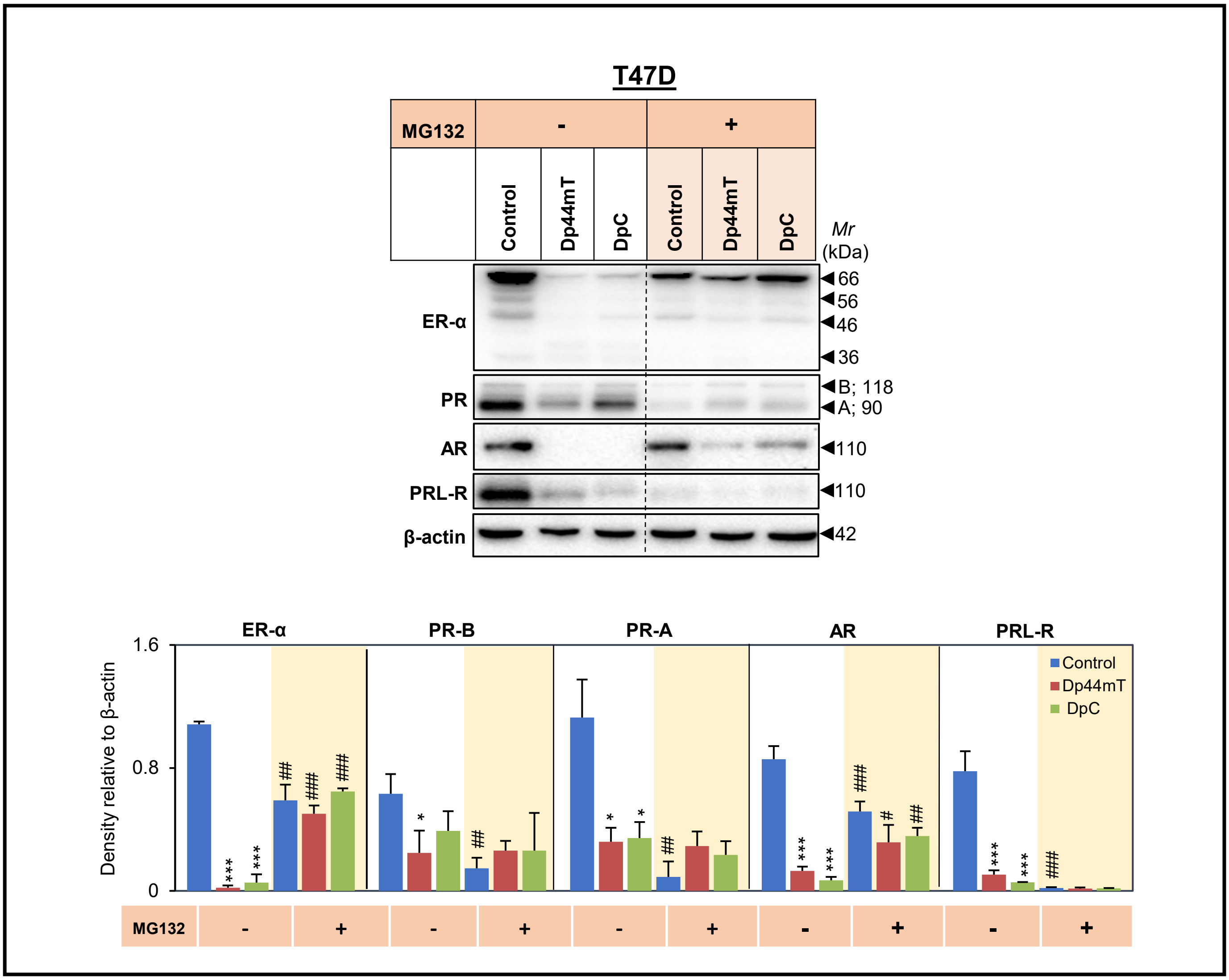


**Supplemental Figure 4**

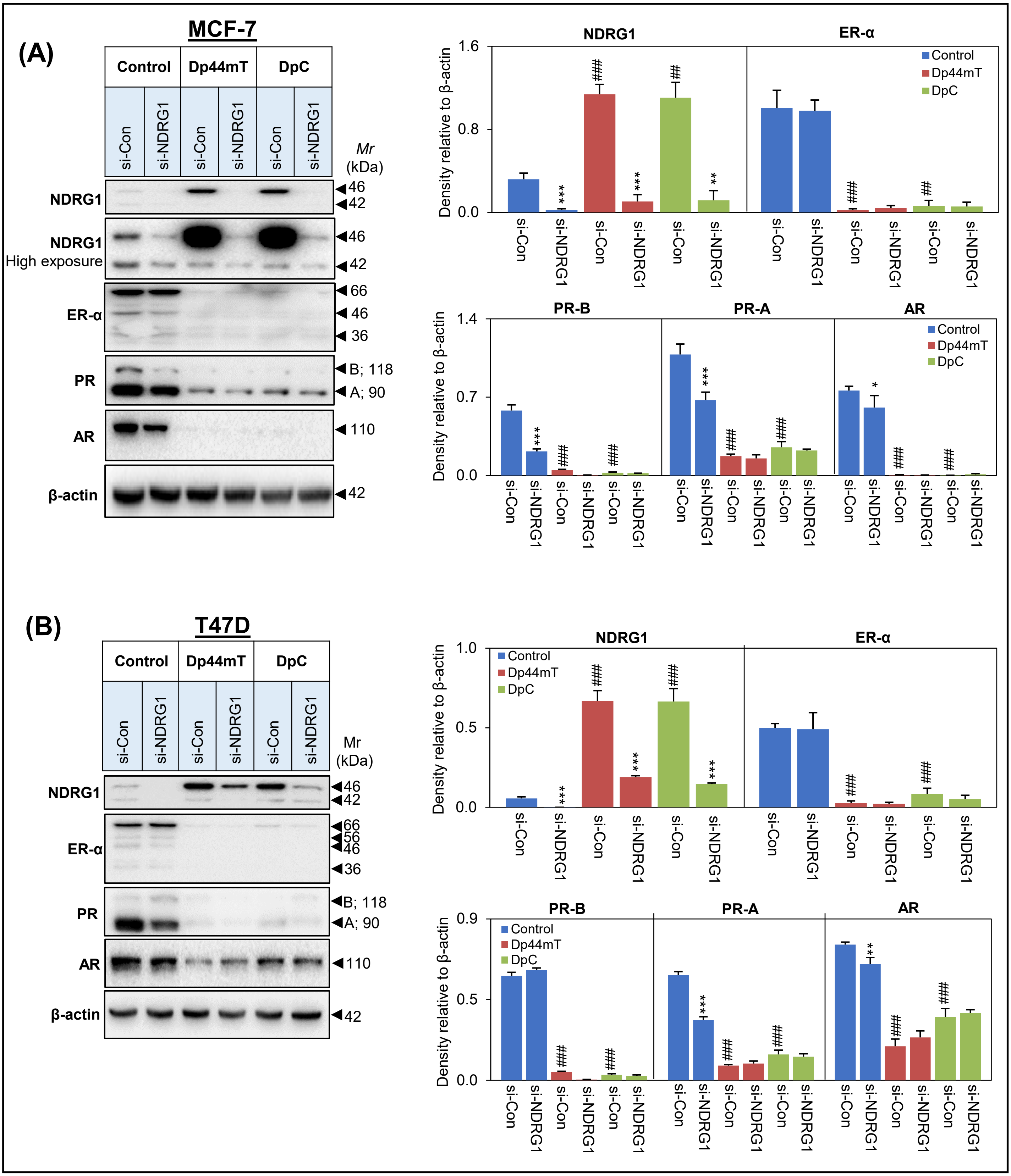


**Supplemental Figure 5**

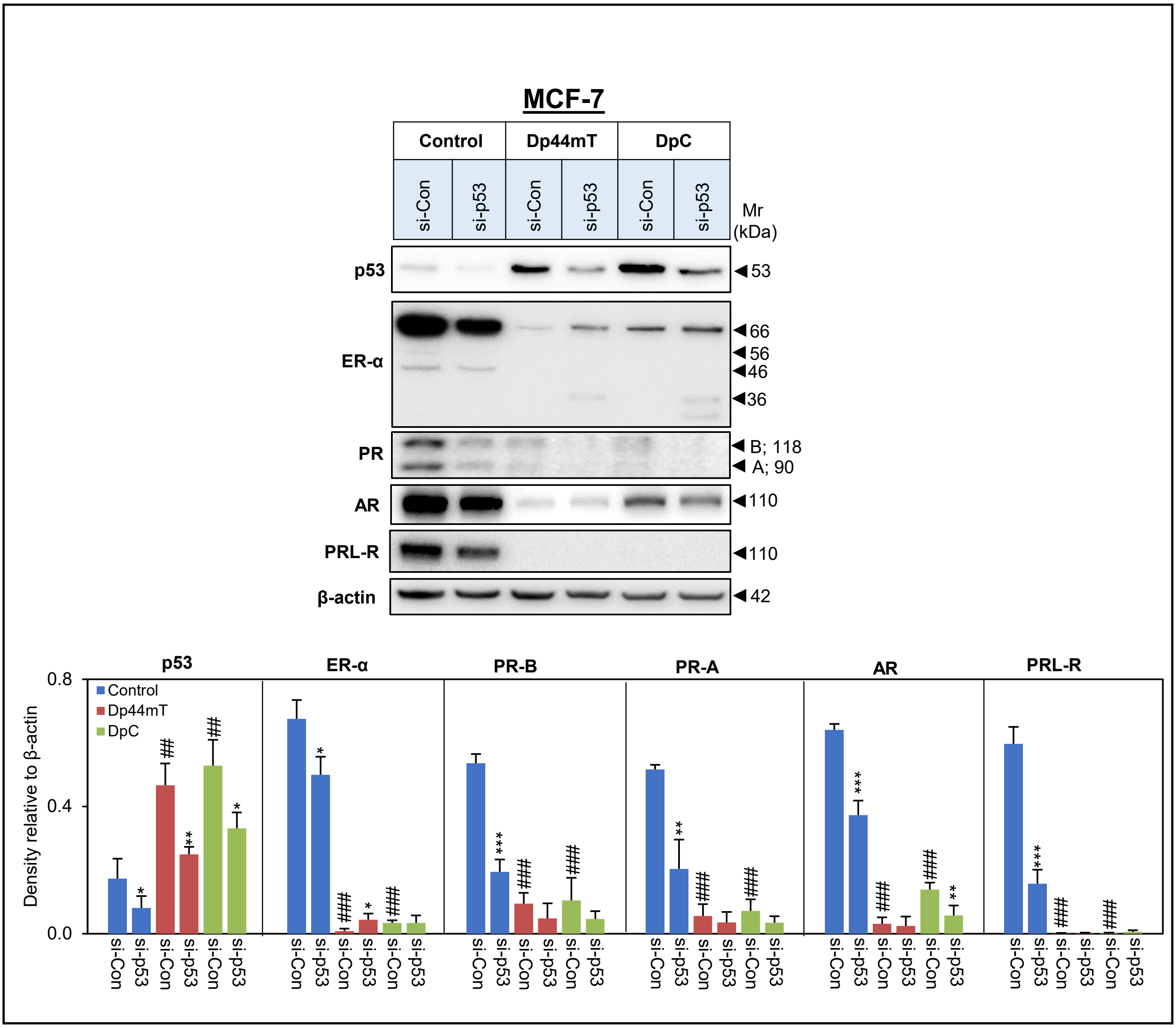


**Supplemental Figure 6**

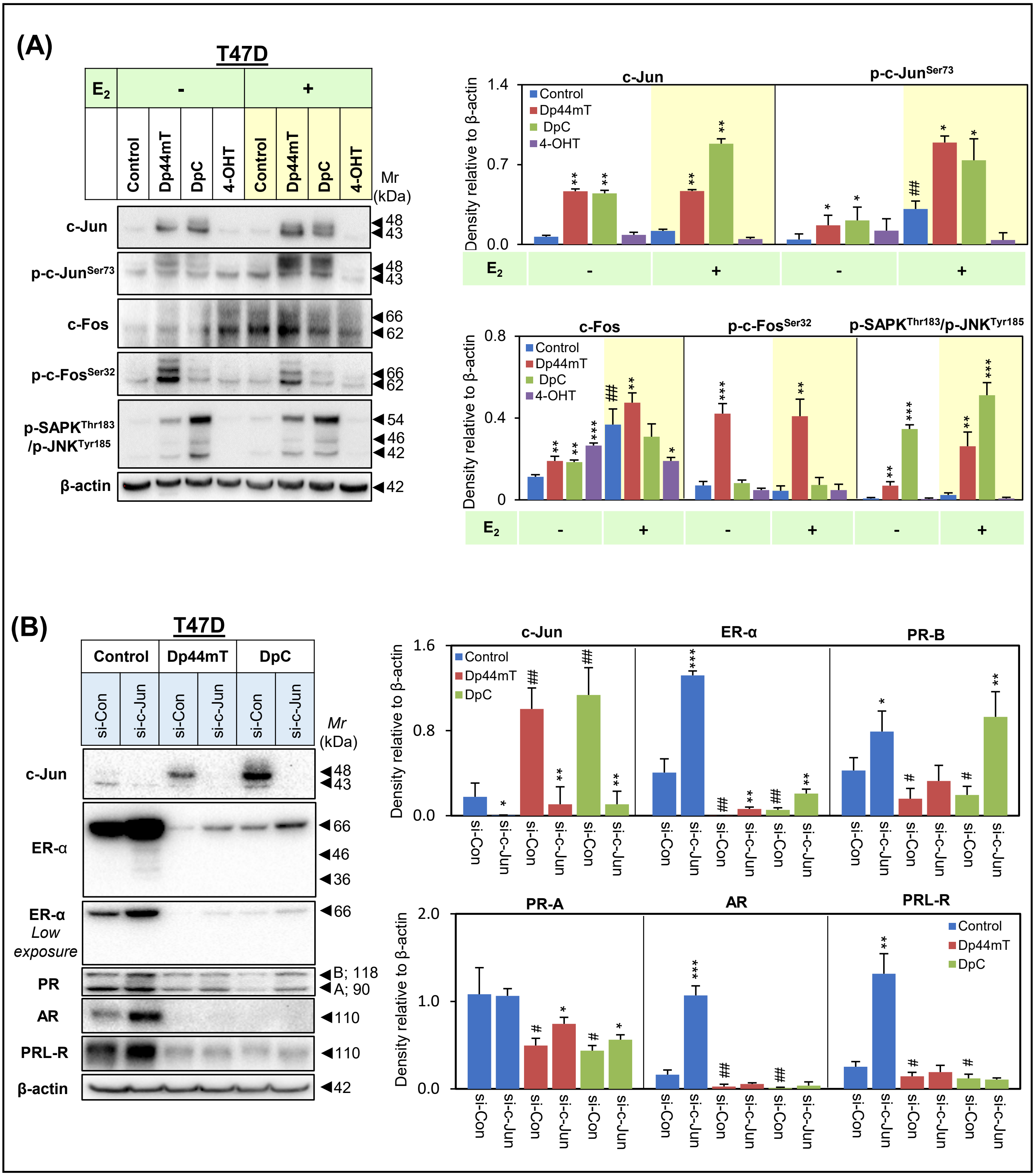


Supplemental Figure 7

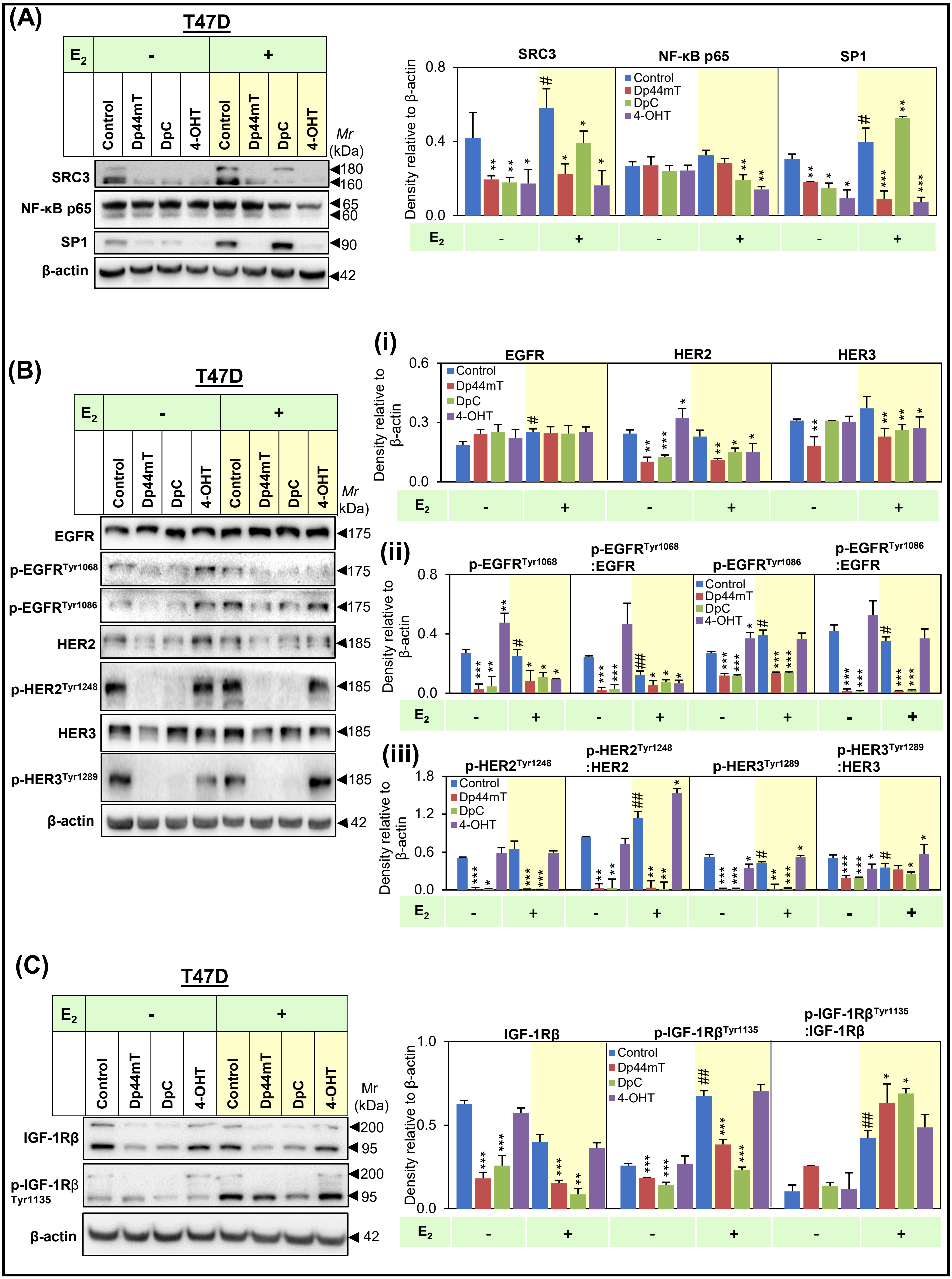


Supplemental Figure 8

**Supplemental Figure Legends**

**Supplemental Figure 1: Dp44mT and DpC down-regulate ER-α, PR, AR, and PRL-R protein levels, but up-regulate NDRG1 protein levels in T47D and ZR 75-1 BC cells. (A)** T47D cells were incubated with control medium or this medium containing Bp2mT (5 µM), DFO (250 µM), Dp44mT (5 µM), DpC (5 µM), or 4-OHT (5 µM) for 24 h. Cellular lysates were then assessed for protein levels of ER-α, PR, AR, PRL-R, and NDRG1 *via* Western blot. **(B)** ZR 75-1 cells were incubated with control medium or this medium containing DFO (250 µM), Dp44mT (5 µM), DpC (5 µM), or 4-OHT (5 µM) for 24 h and then assessed for the protein levels of ER-α, PR, AR, and PRL-R *via* Western blot. β-actin was used as a protein-loading control. Results are mean ± SD ($n = 3$). Significant relative to the untreated control cells: *$p < 0.05$, **$p < 0.01$, ***$p < 0.001$.

**Supplemental Figure 3. Dp44mT and DpC decrease total ER-α and its phosphorylation in T47D BC cells and also MCF-7 BC cells prepared as 3D spheroids. (A)** T47D cells were incubated with control medium or this medium containing Dp44mT (5 µM), DpC (5 µM), or 4-OHT (5 µM) for 24 h, followed by a 15 min incubation in the presence or absence of $E_2$ (10 nM; 15 min). Cellular lysates were then assessed for ER-α, p-ER-$\alpha^{Ser167}$, p-ER-$\alpha^{Ser118}$, PR, AR, and PRL-R *via* Western blot. **(B)** MCF-7 spheroids were grown for 48 h followed by 24 h treatment with control medium, or this medium containing Dp44mT (5 µM) or DpC (5 µM), followed by a 15 min incubation in the presence and absence of $E_2$ (10 nM). Cellular lysates were then assessed for protein levels of ER-α, p-ER-$\alpha^{Ser167}$, and p-ER-$\alpha^{Ser118}$ *via* Western blot. β-actin was used as a protein-loading control. Results are mean ± SD ($n = 3$). Significant relative to the untreated control cells: *$p < 0.05$, **$p < 0.01$, ***$p < 0.001$. Significant relative to $E_2$ untreated cells: $^{\#}p < 0.05$, $^{\#\#}p < 0.01$, $^{\#\#\#}p < 0.001$.

**Supplemental Figure 4. Incubation with DpC decreases protein expression of ER-α, PR, AR, and PRL-R and suppresses their mRNA levels as a function of incubation time in MCF-7 BC cells.**

MCF-7 cells were incubated with DpC (5 μM) for time points ranging from 0 h to 24 h and assessed for: **(A)** Protein levels of ER-α, PR, AR, and PRL-R *via* Western blot; and **(B)** *ER-α*, *PR*, *AR*, and *PRL-R* mRNA levels *via* RT-PCR. β-actin was used as a loading control. Results are mean ± SD ($n = 3$). Significant relative to the untreated (0 h) control cells: $^{**}p < 0.01$, $^{***}p < 0.001$.

**Supplemental Figure 5. Dp44mT and DpC promote proteasomal degradation of ER-α, PR, AR, and PRL-R in T47D BC cells.** T47D cells were incubated with control medium or this medium containing DpC (5 μM) or Dp44mT (5 μM) in the presence or absence of MG132 (2.5 μM). Cellular lysates were then assessed for ER-α, PR, AR, and PRL-R protein levels *via* Western blot. β-actin was used as a protein-loading control. Results are mean ± SD ($n = 3$). Significant relative to untreated control cells: $^{*}p < 0.05$, $^{***}p < 0.001$. Significant relative to corresponding cells in the absence of MG132: $^{\#}p < 0.05$, $^{\#\#}p < 0.01$, $^{\#\#\#}p < 0.001$.

**Supplemental Figure 6. *NDRG1* silencing in MCF-7 and T47D BC cells does not alter the effect of Dp44mT and DpC on decreasing the expression of ER-α, PR, and AR. (A)** MCF-7 and **(B)** T47D BC cells were incubated with negative control siRNA (si-Control) or siRNA against *NDRG1* (si-NDRG1) for 48 h prior to treatment with Dp44mT (5 μM) or DpC (5 μM) for another 24 h. Cellular lysates were then assessed for protein levels of NDRG1, ER-α, PR, and AR *via* Western blot. β-actin was used as a protein-loading control. Results are mean ± SD ($n = 3$). Significant relative to the si-Control cells: $^{*}p < 0.05$, $^{**}p < 0.01$, $^{***}p < 0.001$. Significant relative to the untreated control cells: $^{\#\#}p < 0.01$, $^{\#\#\#}p < 0.001$.

**Supplemental Figure 7. The effect of Dp44mT on decreasing ER-α expression is partly mediated by p53 in MCF-7 BC cells.** MCF-7 cells were incubated with negative control siRNA (si-Control) or siRNA against *p53* (si-p53) for 24 h prior to treatment with either control medium or this medium containing Dp44mT (5 μM) or DpC (5 μM) for another 24 h. Cellular lysates were then assessed for the protein levels of p53, ER-α, PR, AR, and PRL-R *via* Western blot. β-actin was used as a protein-loading control. Results are mean ± SD ($n = 3$). Significant relative to si-Control cells: $^{*}p < 0.05$, $^{**}p < 0.01$, $^{***}p < 0.001$. Significant relative to untreated control cells: $^{\#\#}p < 0.01$, $^{\#\#\#}p < 0.001$.

**Supplemental Figure 8. Dp44mT and DpC promote c-Jun and c-Fos protein levels and activation in T47D BC cells. (A)** T47D cells were incubated with either control medium or this medium containing Dp44mT (5 μM), DpC (5 μM), or 4-OHT (5 μM) for 24 h in the presence or absence of $E_2$ (10 nM). Cellular lysates were then assessed for protein levels of c-Jun, p-c-Jun$^{Ser73}$, c-Fos, p-c-Fos$^{Ser32}$, and p-SAPK$^{Thr183}$/p-JNK$^{Tyr185}$ *via* Western blot. β-actin was used as a protein-loading control. Results are mean ± SD ($n = 3$). Significant relative to untreated control cells: $^{*}p < 0.05$, $^{**}p < 0.01$, $^{***}p < 0.001$. Significant relative to $E_2$ untreated cells: $^{\#\#}p < 0.01$. **(B)** T47D cells were incubated with negative control siRNA (si-Control) or siRNA against *c-Jun* (si-c-Jun) for 24 h prior to treatment with either control medium or this medium containing Dp44mT (5 μM) or DpC (5 μM) for another 24 h. Cellular lysates were then assessed for the protein levels of c-Jun, ER-α, PR, AR, and PRL-R *via* Western blot. β-actin was used as a protein-loading control. Results are mean ± SD ($n = 3$). Significant relative to si-Control cells: $^{*}p < 0.05$, $^{**}p < 0.01$, $^{***}p < 0.001$. Significant relative to untreated control cells: $^{\#}p < 0.05$, $^{\#\#}p < 0.01$.

**Supplemental Figure 9. Dp44mT and DpC decrease protein expression of co-activators, transcription factors, and RTK activation that contributes to ER-α stability and signaling in T47D BC cells.** T47D cells were incubated with control medium or this medium containing Dp44mT (5 μM), DpC (5 μM), or 4-OHT (5 μM) for 24 h in the presence or absence of $E_2$ (10 nM). Cellular lysates were then assessed for protein levels of: **(A)** SRC3, NF-κB p65, and SP1; **(B)** EGFR, p-EGFR$^{Tyr1068}$, p-EGFR$^{Tyr1086}$, HER2, p-HER2$^{Tyr1248}$, HER3, and p-HER3$^{Tyr1289}$; or **(C)** IGF-1Rβ and p-IGF-1Rβ$^{Tyr1135}$. β-actin was used as a protein-loading control. Results are mean ± SD ($n$ = 3). Significant relative to untreated control: $*p < 0.05$, $**p < 0.01$, $***p < 0.001$. Significant relative to cells not treated with $E_2$: $^{\#}p < 0.05$, $^{\#\#}p < 0.01$.
