## Supplemental Table 2 for "An Innovative Non-Hormonal Strategy Targeting Redox Active Metals to Down-Regulate Estrogen-, Progesterone-, Androgen- and Prolactin-Receptors in Breast Cancer"

**Supplemental Table 2:** Differentially regulated genes involved in positive regulation of proteasomal protein catabolic process in response to DpC -treated *vs.* vehicle control-treated MCF-7 cells showing log fold change (logFC), average expression across all samples, in log2 counts per million reads (AveExpr), logFC divided by its standard error (t), Raw p-value (P.Value) and Benjamini-Hochberg false discovery rate adjusted p-value (adj.P.Val).

| ENTREZ ID | SYMBOL | MAP | logFC | AveExpr | t | P.Value | adj.P.Val |
|---|---|---|---|---|---|---|---|
| 9709 | HERPUD1 | 16q13 | 2.74 | 6.78 | 43.33 | 5.78E-12 | 1.67E-09 |
| 127544 | RNF19B | 1p35.1 | 2.65 | 5.92 | 32.69 | 7.00E-11 | 1.06E-08 |
| 3300 | DNAJB2 | 2q35 | 2.23 | 6.82 | 13.26 | 1.83E-07 | 4.22E-06 |
| 54941 | RNF125 | 18q12.1 | 2.08 | 3.37 | 12.84 | 2.40E-07 | 5.30E-06 |
| 1263 | PLK3 | 1p34.1 | 2.99 | 4.09 | 11.16 | 7.86E-07 | 1.42E-05 |
| 10221 | TRIB1 | 8q24.13 | 1.52 | 6.32 | 10.24 | 1.60E-06 | 2.61E-05 |
| 90637 | ZFAND2A | 7p22.3 | 1.45 | 5.36 | 10.00 | 1.95E-06 | 3.09E-05 |
| 1191 | CLU | 8p21.1 | 1.91 | 8.51 | 8.13 | 1.04E-05 | 1.31E-04 |
| 2729 | GCLC | 6p12.1 | 1.85 | 7.70 | 7.90 | 1.30E-05 | 1.59E-04 |
| 57761 | TRIB3 | 20p13 | 1.53 | 6.76 | 7.61 | 1.75E-05 | 2.08E-04 |
| 25897 | RNF19A | 8q22.2 | 1.17 | 7.41 | 3.44 | 3.76E-03 | 2.66E-02 |
